# The evolution of phenotypic plasticity in response to temperature stress

**DOI:** 10.1101/2020.09.15.297515

**Authors:** Francois Mallard, Viola Nolte, Christian Schlötterer

**Author notes:** Corresponding author: Christian Schlötterer.

## Abstract

Phenotypic plasticity is the ability of a single genotype to produce different phenotypes in response to environmental variation. The importance of phenotypic plasticity in natural populations and its contribution to phenotypic evolution during rapid environmental change is widely debated. Here, we show that thermal plasticity of gene expression in natural populations is a key component of its adaptation: evolution to novel thermal environments increases ancestral plasticity rather than mean genetic expression. We determined the evolution of plasticity in gene expression by conducting laboratory natural selection on a *Drosophila simulans* population in hot and cold environments. After more than 60 generations in the hot environment, 325 genes evolved a change in plasticity relative to the natural ancestral population. Plasticity increased in 75% of these genes, which were strongly enriched for several well-defined functional categories (e.g. chitin metabolism, glycolysis and oxidative phosphorylation). Furthermore, we show that plasticity in gene expression of populations exposed to different temperatures is rather similar across species. We conclude that most of the ancestral plasticity can evolve further in more extreme environments.

## Introduction

Phenotypic plasticity is of great interest in ecology and evolution, because it describes the ability of single genotypes to produce distinct phenotypes in different environments (Pigliucci 2001). When populations encounter environmental change, plastic traits will result in phenotypic alterations without genetic response (Price et al. 2003). Of particular importance are those adaptive plastic responses where the altered phenotype is associated with higher fitness, because they provide a selective advantage in variable environments (Charmantier et al. 2008; Dey et al. 2016; Ghalambor et al. 2007; Nussey et al. 2006; Suzuki et al. 2006) or during adaptation to a rapid environmental shift. Phenotypic plasticity is well documented for a broad range of phenotypes including morphological or life history traits (West-Eberhard 2003; Whitman et al. 2009). The technological advances in quantifying gene expression levels for entire transcriptomes have shifted the emphasis to gene expression patterns because many traits/phenotypes can be accurately quantified in a single experiment (Chen et al. 2015; Huang and Agrawal 2016; Zhao et al. 2015).

Despite the conceptual appeal of adaptive plasticity in natural populations, our understanding of phenotypic plasticity in natural populations is still in its infancy (Forsman et al. 2015; Hendry et al. 2016; Merilä et al. 2014; Pigliucci 2005). In addition to adaptive plasticity, traits may be plastic in natural populations for other reasons: 1) neutral plasticity: variation in the trait has no fitness consequences (Via 1993) 2) deleterious plasticity: variation in the expression of the trait may be deleterious and selection operates to minimize it (Dewitt et al. 1998; Ghalambor et al. 2007). The comparison of populations in a common garden experiment is an intuitive and popular approach to infer the selective forces operating on plasticity (Merilä et al. 2014; Levis and Pfenning 2016). Nevertheless, the link between plasticity and adaptation is only correlative and may arise from other changes, not related to adaptation to the environmental contrasts.

Experiments relying on standing genetic variation to study the evolution of plasticity are well-placed in the framework of genetic accommodation (Braendle and Flatt 2006): complex traits with multiple contributing loci can respond quickly to environmental shifts. Hence, phenotypic plasticity could be rapidly modulated in response to selection. Exposing natural populations to more extreme environments provides clear predictions about the evolution of plasticity (Chevin and Hoffmann 2017). While random changes in plasticity are expected under neutrality, in the case of deleterious (costly) plasticity, reduced plasticity is predicted (counter-gradient evolution). An increase in plasticity is expected when plasticity is adaptive: genetic changes in the novel environment will reinforce the ancestral plasticity (Ghalambor et al. 2007, Ho and Zhang 2018). No change in plasticity is difficult to interpret because it may reflect absence of genetic variation, but also weak selection or neutral plasticity result in the same outcome. Experimental evolution is a powerful approach to distinguish between random and directed changes in plasticity because environmental conditions can be tightly controlled and replicated experiments provide more reliable results.

In *Drosophila*, the evolution of gene expression plasticity has been studied for a range of different environmental stressors, ranging from alcohol to heavy metals and temperature (Chen et al. 2015; Huang and Agrawal 2016; Zhao et al. 2015; Clemson et al. 2016; Levine et al. 2011; Porcelli et al. 2016; Yampolski et al. 2012; Zhou et al. 2012). Natural *Drosophila* populations are exposed to daily and seasonal temperature fluctuations (Bergland et al. 2014; Machado et al. 2016), making this a particularly relevant abiotic factor in the context of phenotypic plasticity (Angilletta and Angilletta 2009). Measuring gene expression of a single heterozygous *D. melanogaster* genotype at four different temperatures showed that 83 % of the expressed genes exhibit a plastic expression pattern when exposed to a temperature gradient ranging from 13 to 29 °C (Chen et al. 2015). The variation in gene expression plasticity of natural *Drosophila* populations along latitudinal clines (Zhao et al. 2015; Porcelli et al. 2016) suggests that some of the plastic responses are driven by selection.

We study the evolution of plasticity to infer the influence of high and low temperature regimes on the plasticity of gene expression in *Drosophila simulans* using laboratory natural selection (Fuller et al. 2005, see experimental design in Fig. 1). Specifically, we address the question how adaptation to more extreme temperatures modulates the plastic response of traits, which were already plastic in the founder population. We show that phenotypic plasticity does not prevent evolution. Rather, adaptation to more extreme temperature regimes increases the plastic response. In combination with clinal variation of gene expression in natural populations of both *D. simulans* and *D. melanogaster* (Zhao et al. 2015), our data provide convincing experimental evidence for adaptive phenotypic plasticity in a natural population.

**Fig. 1:**
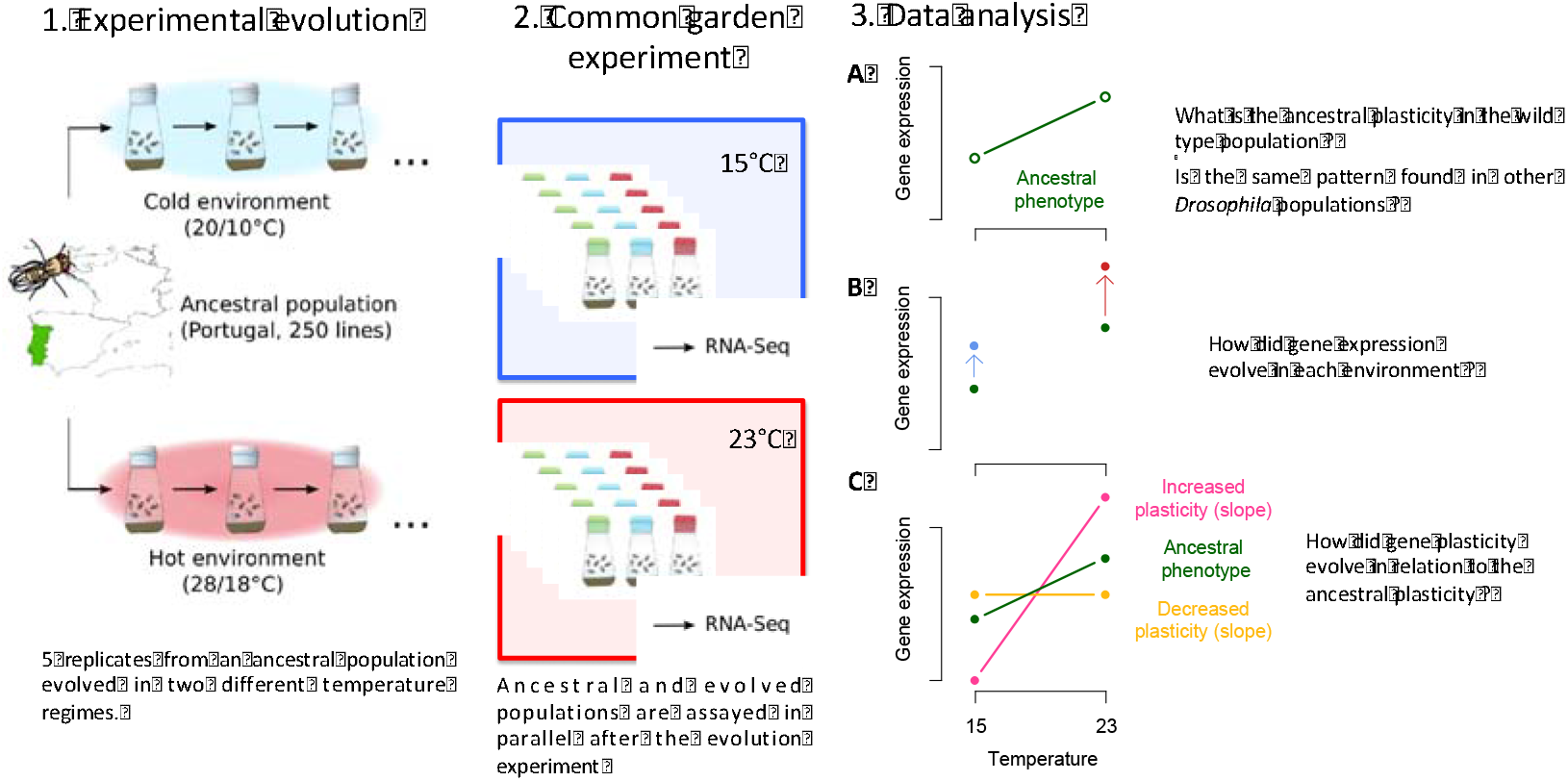
Experimental design. 1) We evolved two sets of five populations in either cold or hot laboratory environments for 39 and 64 generations, respectively. 2) We measured gene expression in two common gardens, where the evolved populations together with the ancestral one were phenotyped at either 15°C or 23°C. 3) Gene expression analysis was done in three successive steps. A) We first explored the plasticity of our ancestral population and compare it to existing data sets. B) We investigated gene expression changes at 15°C and 23°C in the evolved populations. C) We determined the evolved plasticity by measuring each of the evolved populations in both temperature regimes. The evolved plasticity is compared to the ancestral one

## Materials and Methods

### Laboratory natural selection procedure

The laboratory natural selection setup is detailed in Mallard et al. (2018). In brief, 10 replicated *Drosophila simulans* populations were setup from 250 isofemale lines collected in Northern Portugal in 2008. The replicated populations are maintained under two fluctuating temperature regimes (5 replicates in each): either a hot (mean temperature 23°C) or a cold treatment (mean temperature 23°C). In each environment, the temperature changed with a 10°C amplitude centered on the mean temperature synchronized on a 12/12 hours light/dark cycle. The same maintenance regime was used for populations in both temperature environments, only adjusting for the increased developmental time in the cold environment Every generation, 1000 flies are sampled from the eclosed flies and distributed over 5 fresh bottles containing 70 ml standard *Drosophila* medium. After two egg layings for 48h and 72h in the hot and cold environment respectively, adults were frozen. We preferentially used the second egg collection for the next generation to avoid selection for early fecundity. We previously showed that the selection regime results in higher fitness of the evolved populations (Mallard et al. 2018).

### Common garden experiment

Two parallel common gardens with identical experimental procedures were performed in a hot (23°C) and a cold (15°C) environment using eggs from the evolved populations at generation 39 (cold) and 64 (hot). Additionally, 5 replicates of the ancestral population were reconstituted from the founder isofemale lines. After two generations in the assayed environment, the second one with controlled larval density (300 eggs), we collected adults and separated the two sexes under shallow CO_2_. Flies were frozen in liquid nitrogen after a 24h-36h recovery period at 2pm (approximately 6 hours after the start of the light cycle). During experimental evolution, the ancestral population was maintained at 18°C in the form of isofemale lines. The small population size in the isofemale lines prevents adaptation to the culturing conditions and therefore, the reconstituted population reflects the ancestral population (Nouhaud et al. 2016).

### Gene expression analysis

For all 15 populations from both common garden temperatures, we generated two RNA-Seq libraries, each from different sets of 25-30 males. We extracted total RNA-Seq using the Qiagen RNeasy Universal Plus Mini protocol (Qiagen, Hilden, Germany) with DNase I treatment according to the manufacturer’s instructions. Quality control of the RNA was performed on agarose gels and the Qubit RNA HS or BR Assay kit (Invitrogen, Carlsbad, CA) for quantification. Strand-specific barcoded mRNA libraries were generated using the NEBNext^®^ Ultra Directional RNA Library Prep Kit for Illumina with a protocol modified to allow for a larger insert size than the default 200bp. We purified polyA-mRNA from 3μg total RNA and fragmented for 8 min. The 42°C incubation step in the first-strand synthesis and the 16°C step in the second-strand synthesis were extended to 30 and 90 min., respectively. Size selection for a target insert size of 330bp was performed using AMPure XP beads (Beckman Coulter, Carlsbad, CA). PCR amplification followed the recommended protocol (NEB) with 12 PCR cycles and a 50 sec. extension step. The final libraries were bead-purified, quantified with the Qubit DNA HS Assay kit (Invitrogen, Carlsbad, CA) and pooled in equimolar amounts. To reduce batch effects, we combined libraries from ancestral, cold and hot evolved replicates and sequenced them in the same lane. Libraries were sequenced using a single-read 50bp protocol on a HiSeq2500.

We trimmed the raw reads (quality threshold 20, minimum read length 40) using PoPoolation (Kofler et al. 2011). The trimmed reads were aligned to the *Drosophila simulans* reference genome (Palmieri et al. 2014) with GSNAP (Wu et al. 2010) using a hadoop cluster. All subsequent analysis were performed in R (Team RC 2019) including read counts (Liao et al. 2013) and differential gene expression (Robinson et al. 2010). We normalized gene expression levels with the TMM method, restricting our analysis to the genes with an overall mean expression above one count per million (CPM) 11,200 genes). We used negative binomial GLMs to estimate the effect of selection regime, temperature, and their interaction on gene expression. We then computed ad hoc contrasts to find differentially expressed genes between groups of interest using likelihood ratio tests (*glmLRT* in edgeR). This allow us to determine for each gene whether the difference in expression either between two groups of samples (such as the effect of temperature on a given evolved population) or for a linear combination of these groups (such as the difference between the reaction norms of two populations) is statistically significant. The Benjamini-Hochberg procedure was applied to control for false discovery rate (Benjamini et al. 1995). All plasticity estimates as well as evolved differences between ancestral and evolved populations plotted in the manuscript are model fit values obtained from these contrasts.

When comparing the gene expression of evolved populations against the ancestral ones at a given temperature, we always used FDR < 0.05 (unless specified differently). We allowed a higher rate of false positive when testing for reaction norms between ancestral and evolved populations (FDR < 0.1). This was done because we restricted our analysis to genes that were already differentially expressed in at least one temperature with a stringent FDR. Once identified the genes showing a significant evolution of their reaction norms, we compared the absolute value of the ancestral and the evolved reaction norms to distinguish between cases of reduced and increased plasticity. Gene ontology enrichment was performed with Gorilla (Eden et al. 2009) using the complete list of retained genes (n=11,200) as background data set and a FDR<0.05. We compared the number of genes that evolved increased or decreased plasticity in the hot evolved populations using a generalized linear model with a binomial distribution. The estimated probability was compared to the 0.5 using a Wald test.

In a second GLM, we analyzed the replicate specific evolutionary response. We considered only the samples from the Ancestral and the Hot evolved populations and each evolved population was treated as a different level of the “selection regime” factor. The model formula was similar to the previous one but this latter factor contained 6 levels (Ancestral and each of the 5 hot evolved replicate). We processed as described above to detect genes with evolved differential expression.

### Detection of false positive genes with increasing plasticity

To avoid false positives, we restricted our set of candidate genes to those with a significant change in expression in at least one of the two environments (15°C or 23°C) and a significant interaction effect. The rationale can be explained by considering genes that evolved in expression in only one environment, but remained unchanged in the second environment. Adding some minor random noise could either result in a positive or negative correlation of the expression changes in both temperatures. Because negative correlation increases the significance in the interaction test, it may be possible that such random fluctuations could bias our results towards the observed excess of genes with increased plasticity. To rule out that such a potential bias affected our results, we performed an additional test contrasting the ancestral plasticity and the plasticity of a hypothetic population that would have evolved its expression only at one temperature (i.e. replacing the expression levels of the evolved population in the second environment by the ancestral values). For all genes with a significant change in plasticity, we also detected a significant change in plasticity when we considered only the expression change in only one environment. We conclude that none of these genes are false detected due to a random measurement error in the second environment.

### RNA-Seq_quality control

We performed several analyses to test the quality of each library. We first estimated heterogeneity in coverage (3’ bias) of the 20% longest genes of the *D. simulans* annotation using the geneBody coverage tool implemented in the RSeQC package (Wand et al. 2012). Following Mallard et al. (2018), we removed strongly biased libraries (12 libraries in total). Additionally, we quantified the expression of 12 chorion and yolk protein genes to identify female contamination due to sexing mistakes or sample swap. We excluded libraries showing a total log_2_ normalized expression of these genes higher than eight (4 libraries, see Fig. S6). These four libraries contained at least sixteen times the number of transcripts in the remaining libraries (see Fig. S6). After removing the biased and contaminated libraries, a total of 44 libraries remained for the analysis (less than 1.5 samples per population). Out of these 44 libraries, 16 combinations of populations and treatment had only one library left and 14 had 2 libraries. We retained only one measurement per population (n=30) by summing the gene counts of samples coming from the same population. Before pooling the libraries, we visually inspected the samples using multi-dimensional scaling plots (Fig. S1). These plots inform about pairwise distance between samples. While the replicates within a temperature regime were not well separated, robust differences between the ancestral and the two groups of evolved populations were seen. The number of mapped reads for each sample can be found in Table S6.

## Results

We measured the gene expression patterns of our ancestral population and the two evolved populations in two parallel common gardens at 15°C and 23°C. The analysis of evolution of gene expression plasticity is complex and we followed a three-step analysis as described in Figure 1.

### Gene expression plasticity in the ancestral population (Fig. 1 – panel 3A)

We determined temperature mediated plasticity of gene expression by exposing the ancestral, hot evolved and cold evolved population to 15°C and 23°C. As expected from previous studies (Zhou et al. 2012; Chen et al. 2015), the expression of a large number of genes was modulated by temperature.

Down-regulated genes, which are expressed at lower levels at 23°C than at 15°C in the ancestral population, are enriched for several GO categories including chitin-based cuticle and transmembrane transport genes (Table S1). Eighty-nine (83%) of the significant GO terms are also identified among the genes decreasing in expression at higher temperatures in *D. melanogaster* (out of 107 GO terms classified in Chen et al. (2015)). This overlap is probably conservative, because the sex of the flies analyzed and the temperature regimes differed between studies (Chen et al. (2015) measured females in 4 different temperatures). Interestingly Zhao et al. (2015) found that chitin genes were among the top plastic genes shared between *D. melanogaster* and *D. simulans.* In particular, the category “structural constituent of chitin-based cuticle” was consistently identified for genes decreasing with temperature across all three studies.

Genes that are more highly expressed at 23°C than at 15°C in the ancestral population (up-regulated genes) are enriched for genes involved in translation, including a large number of ribosomal genes. Out of 21 GO terms, which were also enriched in Chen et al. (2015), 18 are classified as increasing in both analysis (Table S2). None of these categories were reported in Zhao et al. (2015).

Such highly consistent gene expression changes across different experiments, suggest a highly robust pattern of plasticity, which is conserved not only among populations, but also between species.

### Evolution of gene expression in the focal temperature regime (Fig. 1 - panel 3B)

Only a small number of genes was differentially expressed in populations evolved in the cold environment when compared to the ancestral population (see Table S3 FDR<0.05; 42 genes at 15°C). A quite different pattern was observed for the hot-evolved populations. In the comparison to the ancestral population, 725 genes (see Table S4) were differentially expressed at 23°C. The small impact of adaptation to cold temperature may be the consequence of fewer generations in the new environment compared to the hot-evolved populations. But we cannot distinguish this effect from temperature-specific effects triggering a more pronounced evolution in the hot environment.

### Evolution of gene expression plasticity (Fig. 1 – panel 3C)

With about 32% (n=3,602, FDR<0.05) of the expressed genes being differentially expressed between the two assaying temperatures the cold evolved population was slightly less plastic than the ancestral population (n=4,352, 39%, see Table S1). The hot evolved population had about 44% (n=4,909) plastic genes, which corresponds to about 15% more differentially expressed genes than the other two populations. These differences remain stable even when controlling for the overall library sizes by down sampling (see Fig. S3).

We evaluated the evolution of plasticity by correlating gene expression plasticity (log_2_FC between 15° and 23°C, i.e. the slopes shown in Fig. 1-panel 3C) in the ancestral population with the plasticity in the evolved populations. If the plasticity did not change during evolution, a high correlation is expected. Indeed, the plasticity was highly correlated between ancestral and evolved populations (Pearson correlation coefficients: 0.91 (cold evolved) and 0.89 (hot evolved), Fig. 2). Despite this overall conservation of gene expression plasticity, a closer inspection of Figure 2 (right panel) shows that for some genes plasticity changed after evolution in the hot environment, but the direction of plasticity is not affected (i.e.: the plasticity became more extreme).

**Fig. 2:**
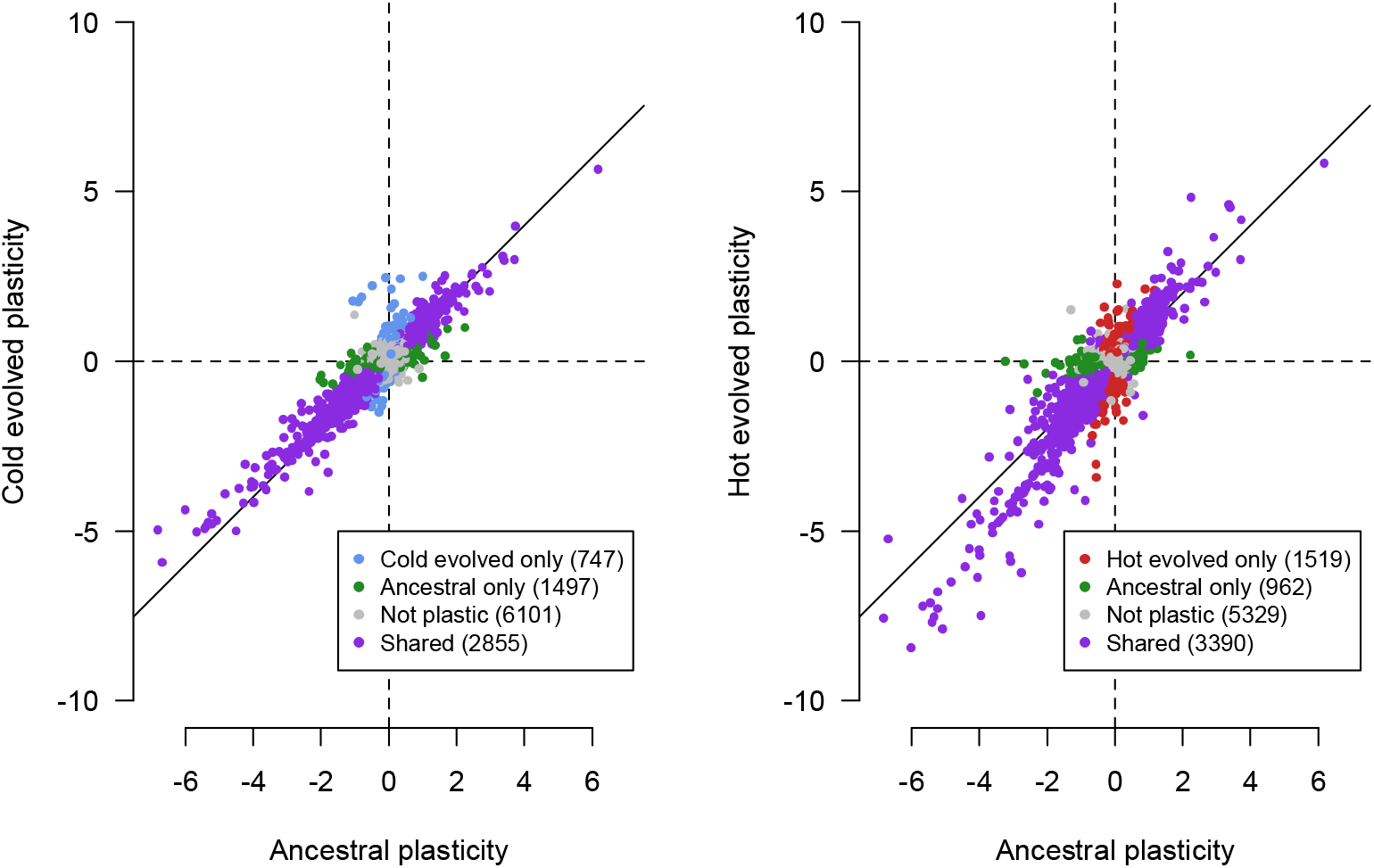
Evolution of gene expression plasticity after selection in cold (left) or hot environments (right). Plasticity is measured as the log_2_ fold change of gene expression at 15°C and 23°C. We compare the plasticity of the ancestral population (x-axis) plotted against the plasticity of the evolved populations. Overall, the pattern of gene expression plasticity is conserved for many genes (purple). Genes that are significantly plastic in only one population have lower log_2_ fold changes in other population (green, blue and red dots). Despite this overall conservation of plasticity, highly plastic genes tend to deviate from the solid line in the hot evolved populations (slope=1) indicating an increased plasticity.

In the cold evolved replicate populations, only a small subset of the genes that evolved a change in expression at 15°C or 23°C displayed a significant difference in the plasticity relative to the ancestral population (2 at 15°C and 2 genes at 23°C, FDR<0.1).

Among the genes that evolved gene expression differences in the hot-evolved populations either at 15°C or at 23°C (n=930), we distinguished three different classes: 1) genes with significant change in plasticity (325 genes, FDR<0.1); 2) genes with small differences in the magnitude of gene expression differences (log_2_FC) between the evolved and the ancestral population in each environment - here, a reliable detection of changes in plasticity or constitutive expression differences is not possible; 3) genes with no change in plasticity, but constitutive expression differences (i.e. a change in the same direction at both temperatures, FDR<0.05, n=50). This third class of genes was enriched for oxido-reduction processes suggesting a global down-regulation of detoxification genes (FDR<0.1, 6 cytochrome p450 genes, 2 UDP-glucuronosyltransferases). Because some of these genes were also down-regulated in the cold evolved populations (23 genes using FDR<0.1 including 6 p450 genes, see Table S3), we conclude that their constitutive change in expression is not directly related to absolute temperature but a response to either temperature stress or to adaptation to shared environmental conditions.

Among the 325 genes with a significant evolution of plasticity, we noticed significantly more genes with increased plasticity (n=241) than with decreased plasticity (n= 84, p<0.001). This result is not biased by ancestrally non-plastic genes that cannot decrease plasticity: the ratio of genes with increased plasticity to genes with decreased plasticity does not change when only ancestrally plastic genes are analyzed (log_2_FC > 1 in the ancestral population, n=62 and 20 respectively, p<0.001). No GO categories were enriched for genes with reduced phenotypic plasticity. In contrast, genes with increased phenotypic plasticity were enriched for several GO terms (116 processes, 34 functions and 28 components). Because the different number of genes in both categories may have affected the enrichment tests, we randomly selected multiple sets of 85 genes among the 242 significant ones and performed the GO analysis for each set. We obtained significantly more enriched processes genes that evolved an increased plasticity (20 bootstrap iterations, p<0.0004; mean number of enriched processes 18.3). Two particularly prominent classes of GO terms were either related to cuticle formation and chitin production or metabolism including the electron transport chain and glucose metabolic processes.

For most of the genes that evolved a difference in gene expression between ancestral and evolved populations, there is a significant change at only one temperature (FDR<0.05). Nevertheless, we noted a strong negative correlation for the sign of the expression differences between hot evolved and founder populations at 15°C and 23°C (Fig. 3, χ^2^_1,241_=133, p<0.0001, see also Fig. S4 for a complementary test), suggesting that evolution modulated the temperature sensitivity of gene expression. This negative correlation is particularly pronounced for genes involved in energy production (see Figs. 4 and S2 for glycolysis and oxidative phosphorylation) but also for chitin related genes.

**Fig. 3:**
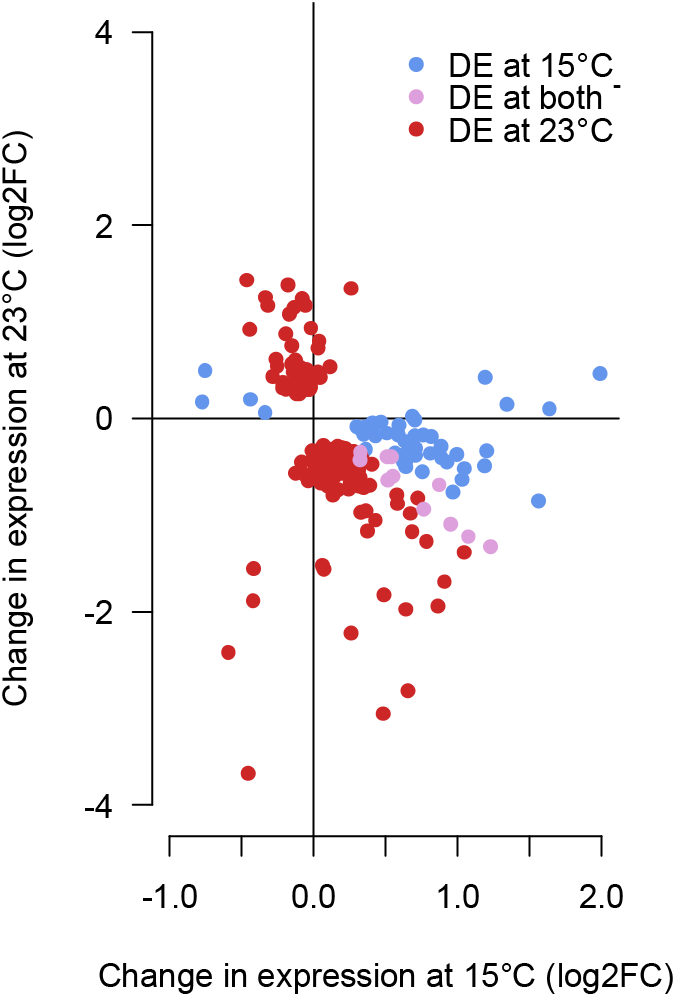
Log_2_FC in gene expression of genes increasing in plasticity during evolution in the hot environment and being differentially expressed in at least one assayed temperature. X- and Y-axis show the impact of adaptation to a hot environment on the gene expression at 15°C and 23°C relative to the ancestral population. Most of these genes evolved a change in expression in the opposite direction (p<0.0001) and are therefore located in the top-left and bottom-right quarters of the plot.

**Fig. 4:**
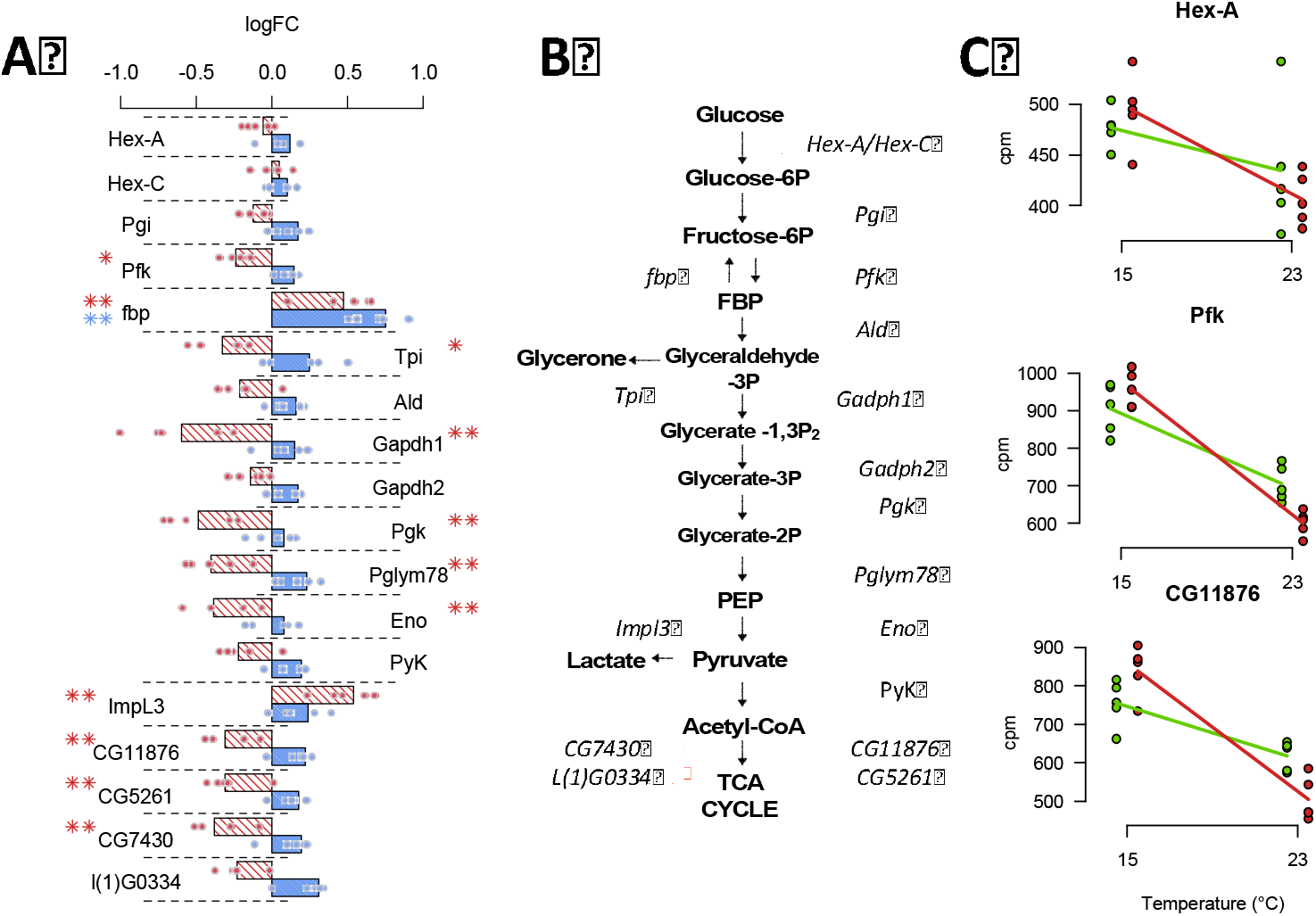
Evolution of plasticity in the glycolysis pathway **A)** Bar chart of the log_2_FC of gene expression evolution from Fig. 2. The difference between the ancestral and the hot evolved populations measured at 15°C (blue) and 23°C (red) are shown. Single dots superimposed on the bar show the divergence of 5 hot-evolved replicates from the mean ancestral expression. **B)** Glycolysis pathway with the main regulatory enzymes from panel A. Most genes involved in glycolysis are significantly down regulated at 23°C (** FDR<0.05, * FDR<0.1). Even for comparisons with no statistically significant difference, most of the genes down regulated at 23°C are up-regulated at 15°C. **C)** Expression plasticity is highly reproducible across replicates. Three enzymes of the glycolysis pathway illustrate the highly consistent response across all five replicates. The ancestral replicate populations are indicated by green dots and the hot evolved populations by red dots (all genes are shown in a Supplementary File). Lines indicate plasticity based on the mean expressions values of the five replicates.

On the other hand, genes that evolved a decreased plasticity show a much weaker correlation of the sign of expression change between temperatures (χ^2^_1,84_=2.8, p=0.09).

### Replicate specific evolution

Our previous analysis is looking for significant changes in expression across five independently evolved populations. Yet, it does not inform us about the parallel evolution of each population. We addressed this by analyzing each replicate independently to detect genes evolving increased or decreased phenotypic plasticity.

In each evolved population, we detected genes that evolved plasticity (range: 85-249, mean =134, 409 genes in total). Only 18 genes were significant in all 5 replicates and 272 in only a single population. Similarly to what we found in the main analysis, more genes displayed increased plasticity (ranging 63-175,n=318) than decreased plasticity (ranging 26-41, n=101). This observation is very consistent across populations: a gene that evolved plasticity in one replicate is found significant in another one with the same frequency (41% and 45% for genes decreasing and increasing plasticity respectively). For changes in reaction norm the consistency across replicates is highly dependent on the direction of change. We observed a low correlation for genes that evolved decreased plasticity (mean r^2^= 0.03, see Fig. S5) while genes that increased plasticity were highly correlated among replicates (mean r^2^=0.77, see Fig. S5).

A GO enrichment analysis at the replicate level showed that only chitin related gene ontologies were significantly enriched in all 5 evolved populations (see Table S5). The increased plasticity of the metabolism related genes were only significantly overrepresented in the first replicate. Nevertheless, we attribute this mainly to a lack of statistical power: the increase in plasticity for the genes involved in glycolysis and oxidative phosphorylation is consistent across all replicates (Fig. 5).

**Fig. 5:**
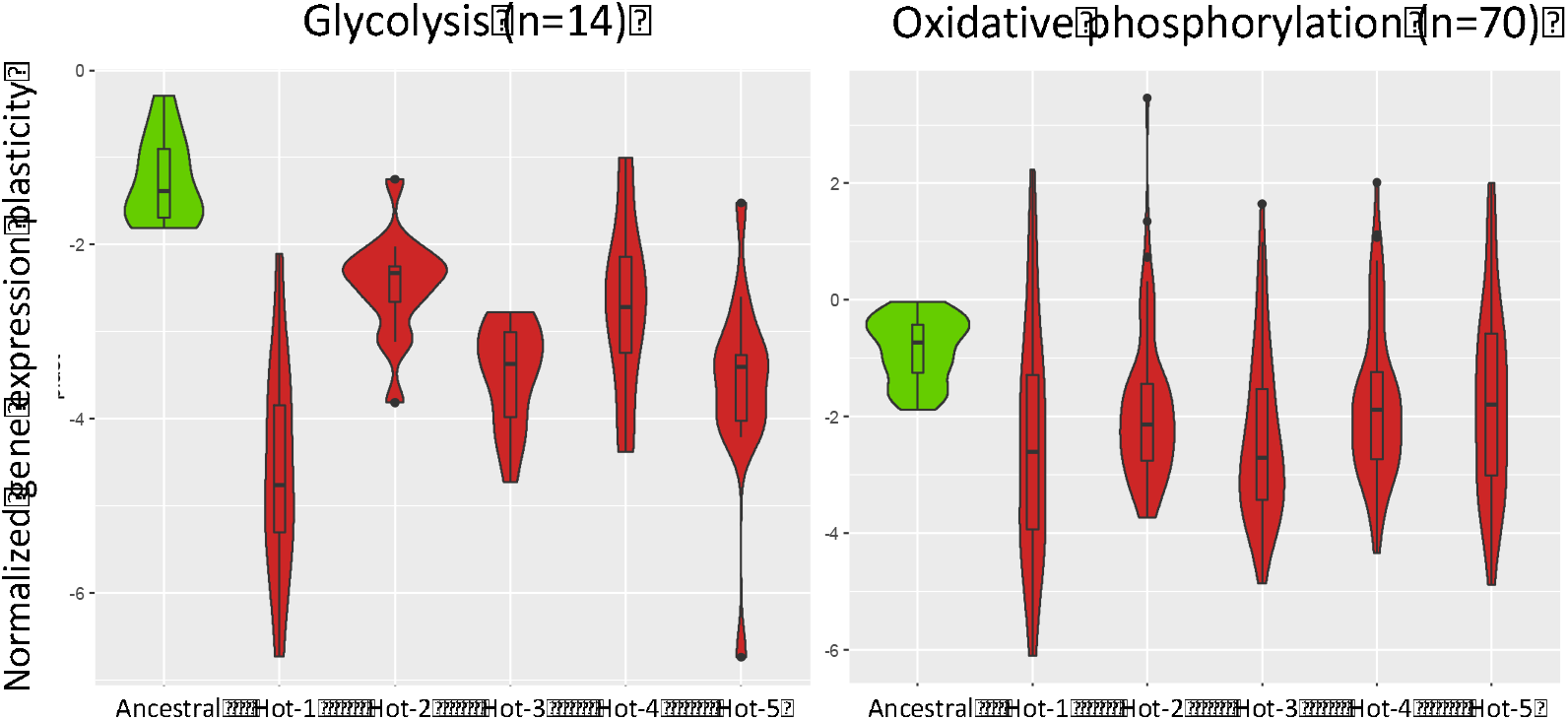
Highly consistent down-regulation of glycolysis and oxidative phosphorylation plasticity across all 5 hot-evolved replicates. The difference in expression between 15°C and 23°C of the ancestral population (green) is higher than in in each of the evolved populations (red) indicating more negative reaction norms. We only show genes from the glycolysis and oxidative pathways that were ancestrally down-regulated (right panel, n=14, left panel, n=70).

## Discussion

### Only limited counter-gradient evolution

Only very few studies were able to address the evolution of gene expression over short evolutionary time scales. The adaptation of *Drosophila melanogaster* to salt and cadmium-enriched medium (Huang and Agrawal 2016) showed that gene expression plasticity evolved, but in the opposite direction to the plasticity seen in the ancestral populations. These food supplements are novel environmental conditions, which are very rarely encountered by fruit flies in their natural environments. The authors proposed that this counter-gradient evolution could be explained by the selection on phenotypes that are only beneficial under these extreme conditions, but not in the environments typically encountered by *Drosophila*: the plastic response would correspond to a “stress” that is no longer expressed when population are adapted to this new environment. This is in sharp contrast to the experimental design of this study. Temperature is one of the most important environmental factors driving local adaptation in ectotherms (Angilletta and Angilletta 2009; Fuller and al. 2005). This applies also to *Drosophila* (Zhao et al. 2015; Bergland et al. 2014; Machado et al. 2016; Klepstatel et al. 2013) where significant clinal variation is seen on the genomic and transcriptomic level (Zhao et al. 2015; Machado et al. 2016; Hoffmann and Weeks 2007). Only a moderate fraction of genes (25% of all genes with evolved plasticity) that experienced counter gradient evolution, i.e. a decrease in the slope of the reaction norm. Interestingly, these genes were not enriched for functional categories and did not evolve consistently across our replicates. Thus, we failed to find biological processes for which gene expression plasticity would be strongly maladaptive. More likely, the gene expression of these genes is not well-adapted, possibly due to pleiotropic functional requirements, which are relaxed in the laboratory environment. The large fraction of genes for which the ancestral plasticity evolved to more extreme values suggests that the laboratory conditions match many ecologically relevant forces encountered by natural *Drosophila* populations.

In our study, we contrasted whole organisms gene expression across environments - a common practice in the study of gene expression evolution in *Drosophila* (Chen et al. 2015a, 2015b; Huang and Agrawal 2016; Zhao et al. 2015; Clemson et al. 2016; Levine et al. 2011; Porcelli et al. 2016; Yampolski et al. 2012; Zhou et al. 2012). Nevertheless, a potential problem is that during evolution allometric changes may occur - this is that the relative abundance of some cell types changes (Montgomery and Mank 2016). In fact, a recent study showed that females adapting to a new temperature regimes also evolved allometric changes, while males were much less affected (Hsu et al. 2020). While such allometric changes could affect gene expression means, the impact on plasticity is not clear. If the evolved allometric changes do not change with assaying temperature, no influence on the analysis of phenotypic plasticity is expected. On the other hand, if allometric changes are modulated by assaying temperature, this could be considered as an extended evolved phenotypic plasticity and will not affect our conclusions.

### Plasticity in gene expression suggests adaptive plasticity

The evolutionary implication of phenotypic plasticity is a controversial topic with two extreme perspectives. With the same genotype expressing different phenotypes in response to the environment, it is often assumed that these phenotypes provide a higher fitness to their carriers (Via et al. 1995). If phenotypic plasticity results in a good match of phenotype and environment, this could even make genetic adaptation expendable (Charmantier et al. 2008). On the other hand, phenotypic plasticity of many traits may not contribute to fitness and reflects pleiotropic responses to environmental changes. This uncertainty about the evolutionary consequences has not yet been settled because of the difficulty to link plasticity with fitness advantage. Our study links the evolutionary response in a laboratory natural selection experiment to plasticity in the founder population. Out of 3,605 genes with plastic gene expression pattern after exposure to two temperatures, 327 genes (9%) changed plasticity after 59 generations. Reasoning that the hot laboratory environment is more extreme than the habitat of the founder population, genes with adaptive plasticity for temperature should evolve towards increased plasticity (Lande 2009; Garland and Kelly 2006). Consistent, with this expectation, 75% of the genes with evolved plasticity increased their environmental sensitivity. Genes with increased thermal sensitivity showed functional enrichment and were more consistent in their change across replicates. Our gene expression results are in line with the prevalence of genetic variation for thermal plasticity in natural *Drosophila* populations (Zhao et al. 2015; Levine et al. 2011, but see Clemson et al. 2016). The parallel evolution in plasticity suggests a selective advantage of populations with evolved plasticity, but our experiment cannot decide whether the evolved plasticity is providing the fitness advantage or it is a pleiotropic effect caused by the true target of selection. Finally, most of the genes with increased plasticity showed an opposite evolutionary response at 15°C and 23°C leading to the reinforcement of the ancestral plasticity which is expected in the case of adaptive plasticity (Ghalambor et al. 2007, Ho and Zhang 2018).

Future experiments, measuring individual flies gene expression would allow us to study the evolution of the trait gene expression plasticity. Comparing the trait distribution in the ancestral and evolved populations after the new trait optimum has been reached will provide further insights in the underlying adaptive architecture.

### Evolution of plasticity is more frequent than constitutive expression changes

In hot evolved populations, only 7% (52 out of 729) of the genes, which evolved a significant response relative to the founder population at 23°C, showed a constitutive expression difference rather than an evolutionary change of plasticity. It is not clear if this predominance of plastic response reflects the design of the laboratory natural selection experiment, which involved daily temperature fluctuation or a correlated response to directional selection (Garland and Kelly 2006).

While the flies evolved in a novel temperature regime with daily fluctuations, we measured gene expression in constant temperature regimes to avoid confounding effect of development at different temperatures. Classic examples for the persisting effects of short-term exposure to high temperatures are phenocopies. Short (<5h) sensitive periods of *Drosophila* pupae result in different phenotypes depending on the developmental stage during exposure (Mitchell and Lipps 1978). Hence, even small differences in developmental timing could result in large phenotypic variation within or between populations. Thus, we opted for a constant temperature common garden. This strategy assured phenotypic measurements insensitive to daily temperature fluctuations, reflecting fixed temperature effects that are comparable to existing phenotype data. Given that the expression of most genes changes monotonically with temperature (Chen et al. 2015a), we anticipated that observed differences in reaction norm at 15 and 23°C can be extrapolated to more extreme temperatures, such as 10 and 28°C.

While it is possible that the observed gene expression changes are not the direct target of selection, it would not challenge our claim that ancestral plasticity is likely to be adaptive: even if the evolution of the gene expression in our experiment is only correlated with the selected trait(s), the ancestral plasticity we observed at the gene expression level remains an indicator of adaptive plasticity because the direction of change is, by definition, the same between these correlated traits. The validity of our conclusions would only be challenged if during evolution these phenotypic correlations across temperature were broken. We consider this, however, unlikely as the temperature response is conserved across populations and various *Drosophila* species (see also Zhao et al. 2015).

We previously identified SNF4Aγ and Sestrin as targets of selection in the same hot evolved populations (Mallard et al. 2018). Both genes are associated with activity of AMPK, a key enzyme in metabolism regulation. Interestingly, the role of AMPK in thermal plasticity have been highlighted in marine invertebrates such as mussels and rock crabs that are regularly subjected to temperature variation during tides (Frederich et al. 2009; Jost et al. 2014). Moreover, mussels experience seasonal variation in thermal plasticity of AMPK activity (Jost et al. 2014), which is comparable to the evolution of plasticity in our evolved populations. In addition to metabolism, chitin synthesis was found to be plastic, which is shared with *D. melanogaster* (Chen et al. 2015) and in the North American cline (both *D. melanogaster* and *D. simulans* (Zhao et al. 2015)). Chitin is involved in exoskeleton morphogenesis and its decreased synthesis may be associated with the temperature-induced size reduction in *Drosophila*. However, we did not find any evolution of body size during our experiment (data not shown). Alternatively, chitin is also essential for trachea formation (Moussian et al. 2005), and the evolution of its synthesis in our experiment could be linked with the decrease in metabolism gene expression.

Previous experimental evolution studies in *Drosophila* have found inconsistent results regarding the evolution of gene expression plasticity (Huang and Agrawal 2016; Yampolski et al. 2012) and it is not clear if this inconsistency can be explained by different environmental stressors. On the other hand, it has been proposed that plasticity increases during the initial phase of adaptation to novel environments, followed by genetic assimilation (Lande 2009). In this theoretical scenario, also called “plasticity first” (Levis and Pfenning 2016; Levis et al. 2018), the genomic variation which encodes phenotypic plasticity is favored as a rapid phenotypic response. As a consequence, selection signatures are expected for genes modulating plasticity, rather than in cis-regulatory variation of genes with modified gene expression patterns.

In the context of the current ongoing climate change, the role of phenotypic plasticity has been widely discussed - does plasticity favor or limit genetic adaptation (Merilä and Hendry 2014; Sgrò et al. 2016; Vazquez et al. 2017; DeBiasse et al. 2018)? As recently stated by Kelly (2019), if plasticity is a major contributor of adaptation to climate change, then the amount of available genetic variation for plasticity could be a reliable predictor of a population vulnerability. In particular, for *Drosophila*, the potential of plasticity for attenuating the impact of climate change has been challenged. Thermal plasticity does not correlate with latitude (Sørensen et al. 2016) and did not respond to laboratory natural selection in higher order phenotypes when submitted to stable or fluctuating environments (Fragata et al. 2016; Manenti et al. 2015). Our experiments provide some important insights into this debate. The highly parallel response in replicated populations demonstrates that genetic variation in thermal plasticity is a reservoir for adaptation in novel thermal environments.

Because we studied plasticity after only a moderate number of generations, our study is not informative for more long-term evolutionary processes. Recently, it has been shown that on the long term this evolutionary response could lead to extinction unless a small number of genetic loci are involved (Nunney 2016). Whether a phase of genetic assimilation will follow this initial increase in plasticity will depend on the availability of the relevant variation. If such variants are still segregating, it could be informative to test our experimental populations at later generations. If new mutations are required, experimental evolution in *Drosophila* may not be well-suited to address this question because the spread of new mutations is rare (Burke et al. 2010).

## Data Accessibility

Sequence reads from this study will be available at the European Sequence Read Archive (http://www.ebi.ac.uk/ena/). Additionally, we will provide our gene expression count table and all our R code on dryad.

## Supplemental Figures

**Fig. S1:**
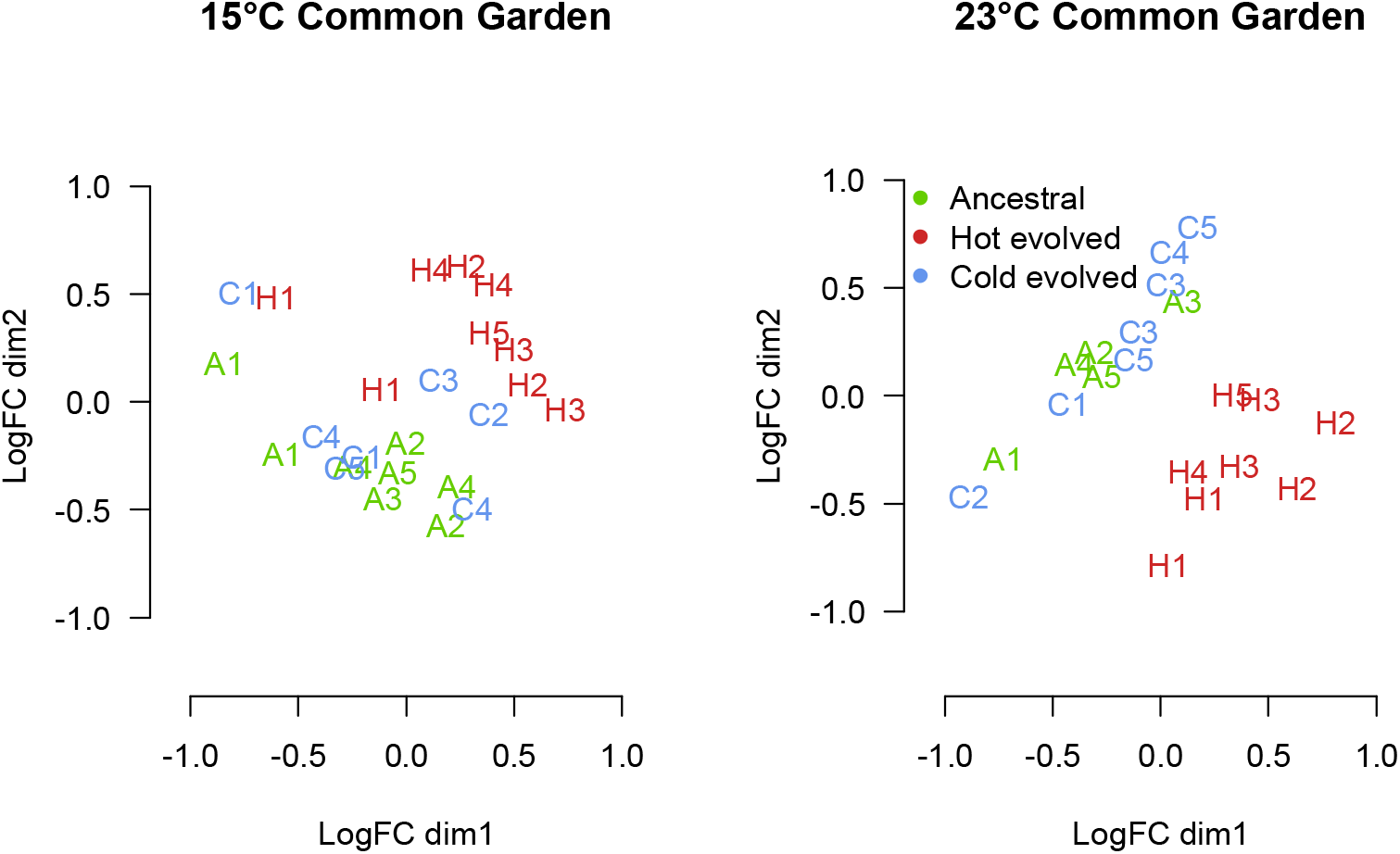
Multi-dimensional scaling plots of the different samples. Ancestral, Cold evolved and Hot evolved population are respectively labeled A[1-5], C[1-5] and H[1-5] and colored in green, blue and red. Left and right panels respectively show the samples of the 15°C and 23°C common garden experiments. In both experiments, we did not detect any outliers.

**Fig. S2:**
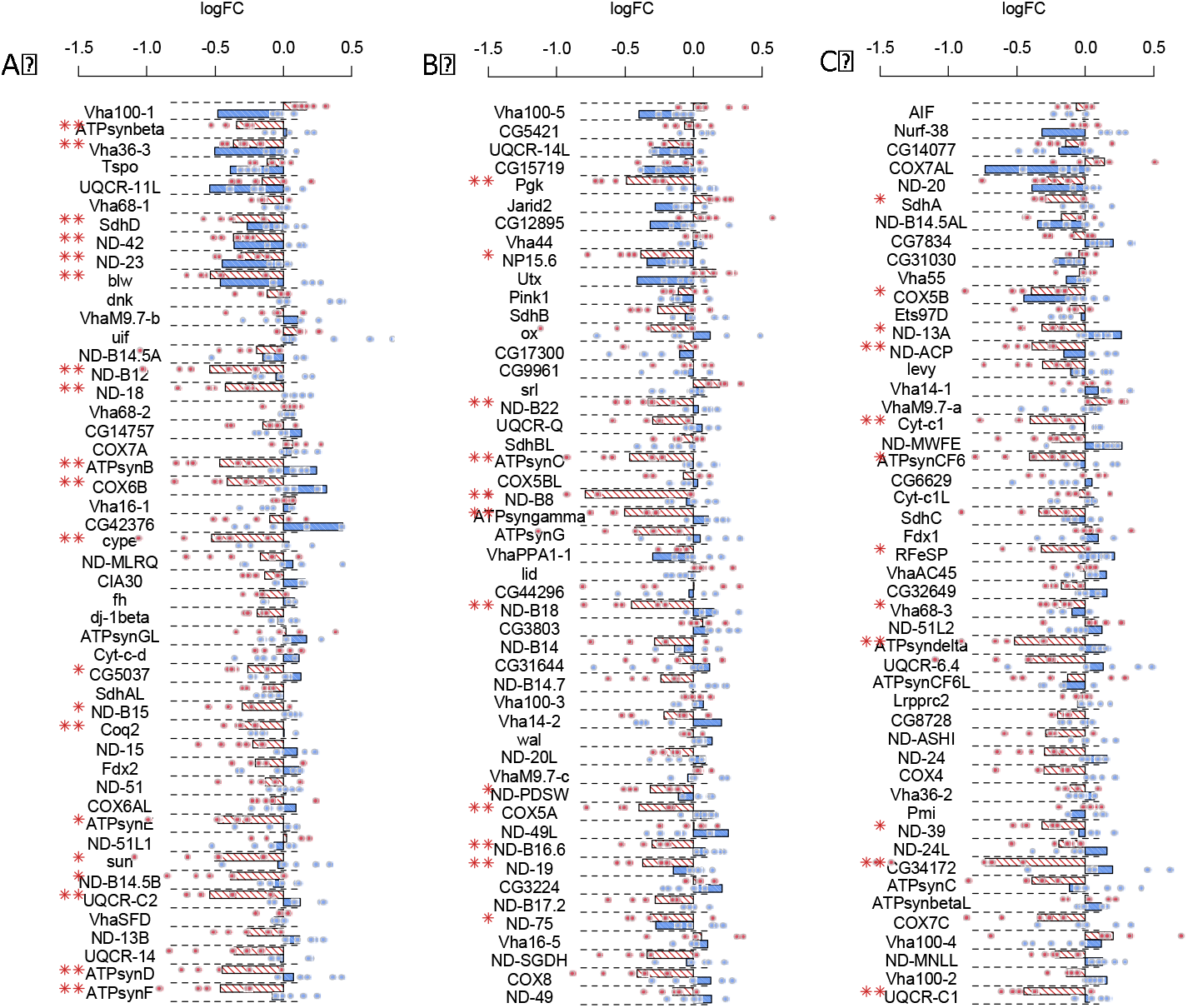
Log_2_FC of gene expression between the ancestral and the hot evolved populations at 15°C (blue) and 23°C (red) of the 146 genes involved in the oxidative phosphorylation. Most of these genes are significantly down regulated at 23°C (n=45, ** FDR<0.05, * FDR<0.1). Even though most comparison are not statistically significant, most of the down regulated genes at 23°C are up-regulated at 15°C. We produced independent plot for each gene showing the variability across our five replicates in a Supplementary File.

**Fig. S3:**
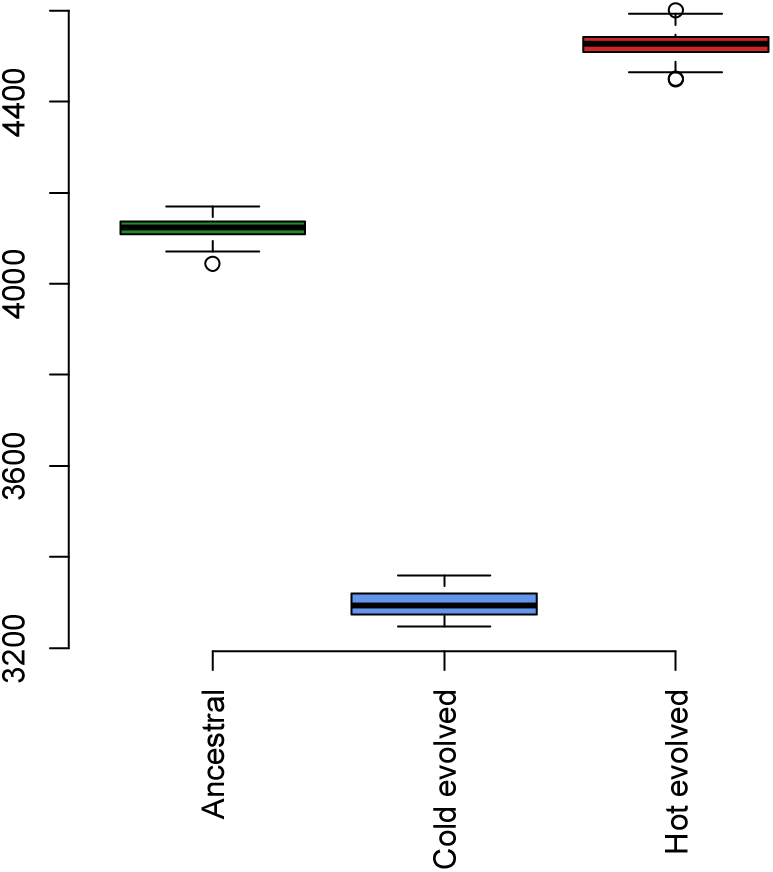
The number of genes statistically plastic in the ancestral and the two groups of evolved populations after down-sampling each library to the same number of reads (14,464,603). These distribution result from 100 independent sampling of the complete libraries. Although the total number of plastic genes is smaller after the down-sampling, the differences between groups are similar.

**Fig. S4:**
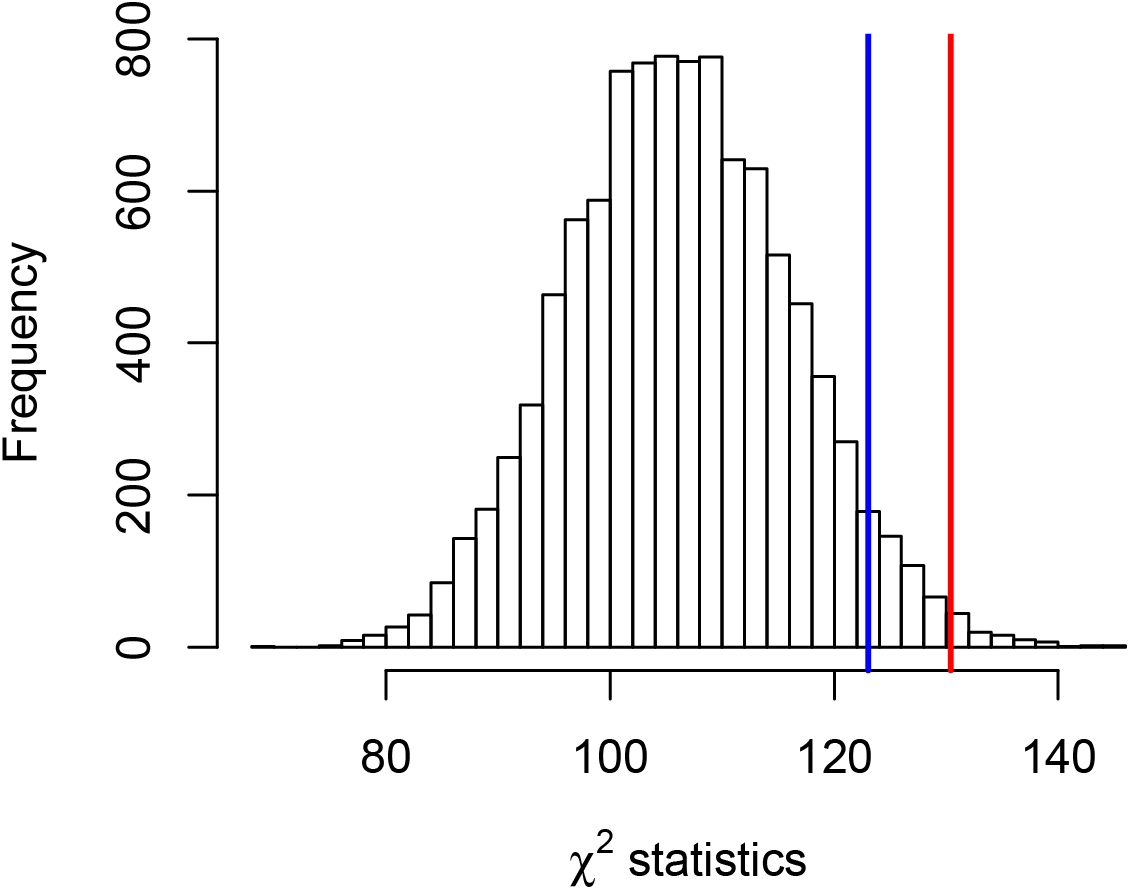
The direction of the evolved gene expression change is highly correlated at 15°C and 23°C. Because the genes are unlikely to evolve independently, the χ^2^ assumptions are violated. To overcome this limitation, we generated an empirical distribution of χ^2^ statistics by bootstrapping the genes that had a significant change in plasticity during evolution in the hot environment and still found a more significant correlation for the sign of gene expression change at 15°C and 23°C than expected by chance. This empirical distribution of χ^2^ statistics was obtained by bootstrapping 241 genes 10,000 times from the list of 417 genes that evolved a significant change in plasticity (FDR<0.1, here we did not condition on a significant evolution at 15°C or 23°C). For each list of 241 genes, a χ^2^ test of independence was computed between the direction of the evolved expression change at 15°C and 23°C. The statistic obtained using the genes that evolved increased plasticity (observed data, red line) is larger than 99% of the statistics obtained by bootstrapping (the blue line shows the 95% threshold). This test indicates that the negative correlation observed for the list of genes showing increasing plasticity could not be obtained from a random sampling of genes showing overall plasticity evolution. Here we assume that both lists of genes have equal modularity: the evolution of a subset of these genes is due to a similar number of causative events.

**Fig. S5:**
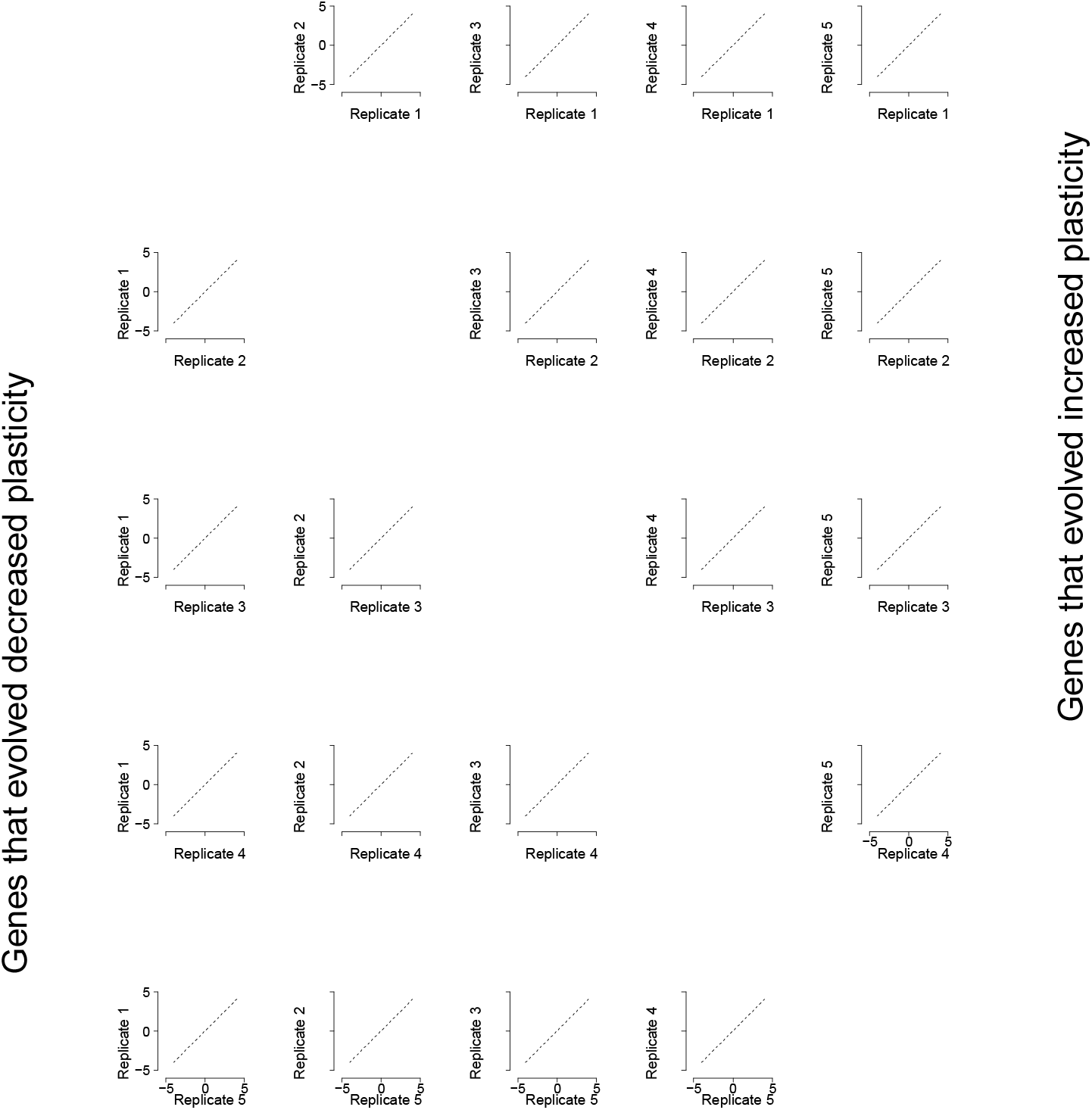
Pairwise log_2_FC of reaction norm slopes between five evolved replicates (as rows and columns). The bottom half displays the genes with decreased plasticity during evolution and the top half the genes with increased plasticity during evolution (see Fig 1-3C with matching color code). There is a strong correlation of the plasticity evolution across all 5 hot-evolved replicates for genes with increased plasticity (top) but not for genes with decreased plasticity (bottom).

**Fig. S6:**
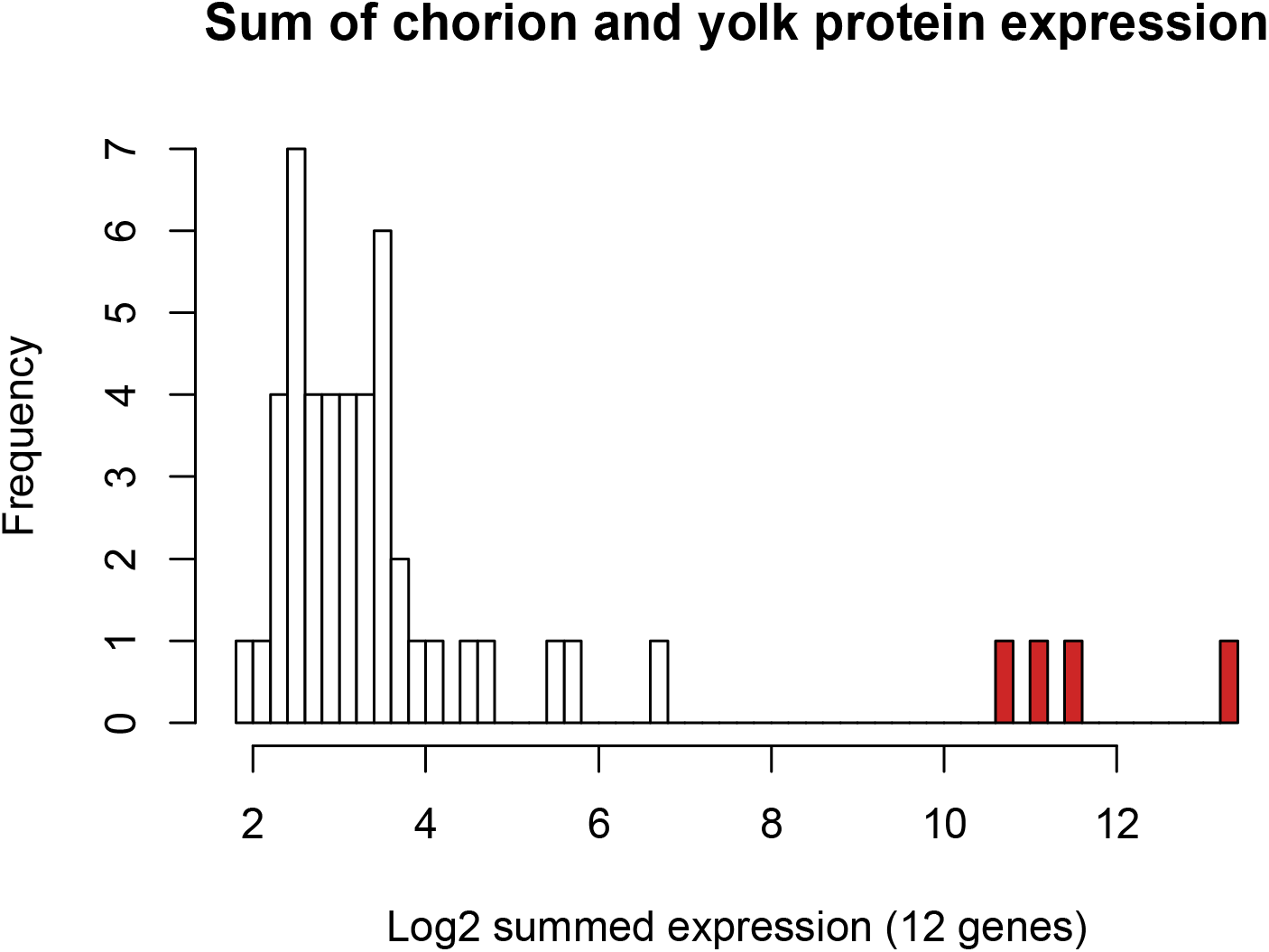
Identification of libraries with female contamination. We summed the expression of the nine chorion genes (CP15 to 19, CP36, CP38) and three yolk proteins (YP1 to 3). We excluded four outlier libraries (in red) with > 1769 counts per million reads for the 12 indicator genes. The retained libraries had < 111 counts per million reads.

## References

Angilletta Jr MJ, and Angilletta MJ. Thermal adaptation: a theoretical and empirical synthesis. Thermal adaptation: a theoretical and empirical synthesis. Oxford University Press; 2009.

Angilletta MJ, and Dunham AE. The temperature-size rule in ectotherms: simple evolutionary explanations may not be general. Am Nat United States; 2003;162:332–42.

Benjamini Y, and Hochberg Y. Controlling the false discovery rate: a practical and powerful approach to multiple testing. Journal of the royal statistical society. Series B (Methodological). JSTOR; 1995;:289–300.

Bergland AO, Behrman EL, O’Brien KR, Schmidt PS, and Petrov DA. Genomic evidence of rapid and stable adaptive oscillations over seasonal time scales in Drosophila. PLoS Genetics. United States; 2014;10:e1004775.

Braendle C, and Flatt T. A role for genetic accommodation in evolution? Bioessays. United States; 2006;28:868–73.

Burke MK, Dunham JP, Shahrestani P, Thornton KR, Rose MR, and Long AD. Genome-wide analysis of a long-term evolution experiment with Drosophila. Nature. England; 2010;467:587–90.

Charmantier A, McCleery RH, Cole LR, Perrins C, Kruuk LEB, and Sheldon BC. Adaptive Phenotypic Plasticity in Response to Climate Change in a Wild Bird Population. Science. 2008;320:800–803.

Chen J, Nolte V, and Schlötterer C. Temperature stress mediates decanalization and dominance of gene expression in Drosophila melanogaster. PLoS Genet. Public Library of Science; 2015;11:e1004883.

Chen J, Nolte V, and Schlötterer C. Temperature-Related Reaction Norms of Gene Expression: Regulatory Architecture and Functional Implications. Molecular Biology and Evolution. United States; 2015;32:2393–402.

Chevin L-M, and Hoffmann AA. Evolution of phenotypic plasticity in extreme environments. Philosophical Transactions of the Royal Society of London B: Biological Sciences. The Royal Society; 2017;372.

Clemson AS, Sgrò CM, and Telonis-Scott M. Thermal plasticity in Drosophila melanogaster populations from eastern Australia: quantitative traits to transcripts. J Evol Biol. Switzerland; 2016;29:2447–2463.

DeBiasse, MB, Kawji, Y, Kelly, MW. Phenotypic and transcriptomic responses to salinity stress across genetically and geographically divergent *Tigriopus californicus* populations. Mol Ecol. 2018;27:1621–1632.

Dewitt TJ, Sih A, and Wilson DS. Costs and limits of phenotypic plasticity. Trends Ecol Evol. England; 1998;13:77–81.

Dey S, Proulx SR, and Teotónio H. Adaptation to Temporally Fluctuating Environments by the Evolution of Maternal Effects. PLoS Biol. 2016;14:e1002388.

Eden E, Navon R, Steinfeld I, Lipson D, and Yakhini Z. GOrilla: a tool for discovery and visualization of enriched GO terms in ranked gene lists. BMC bioinformatics. BioMed Central; 2009;10:1.

Forsman A. Rethinking phenotypic plasticity and its consequences for individuals, populations and species. Heredity (Edinb). England; 2015;115:276–84.

Fragata I, Lopes-Cunha M, Bárbaro M, Kellen B, Lima M, Faria GS, Seabra SG, Santos M, Simões P, and Matos M. Keeping your options open: Maintenance of thermal plasticity during adaptation to a stable environment. Evolution. United States; 2016;70:195–206.

Franssen SU, Kofler R, Schlötterer C. Uncovering the genetic signature of quantitative trait evolution with replicated time series data. Heredity. 2017;118:42–51.

Fuller RC, Baer CF, and Travis J. How and When Selection Experiments Might Actually be Useful. Integrative and Comparative Biology. Oxford University Press; 2005;45:391–404.

Garland T, and Kelly SA. Phenotypic plasticity and experimental evolution. J Exp Biol. England; 2006;209:2344–61.

Ghalambor C., McKay J., Carroll S., and Reznick D.. Adaptive versus non-adaptive phenotypic plasticity and the potential for contemporary adaptation in new environments. 2007.

Ghalambor CK, Hoke KL, Ruell EW, Fischer EK, Reznick DN, and Hughes KA. Non-adaptive plasticity potentiates rapid adaptive evolution of gene expression in nature. Nature. England; 2015;525:372–5.

Hendry AP. Key Questions on the Role of Phenotypic Plasticity in Eco-Evolutionary Dynamics. J Hered. United States; 2016;107:25–41.

Ho WC, and Zhang J. Evolutionary adaptations to new environments generally reverse plastic phenotypic changes. Nat Commun. England; 2018;9:350.

Hoffmann AA, and Weeks AR. Climatic selection on genes and traits after a 100 year-old invasion: a critical look at the temperate-tropical clines in Drosophila melanogaster from eastern Australia. Genetica. Netherlands; 2007;129:133–47.

Hsu, S. K., Jakšić, A. M., Nolte, V., Lirakis, M., Kofler, R., Barghi, N., … & Schlötterer, C. (2020). Rapid sex-specific adaptation to high temperature in Drosophila. eLife, 9, e53237.

Huang Y, and Agrawal AF. Experimental Evolution of Gene Expression and Plasticity in Alternative Selective Regimes. PLOS Genetics. 2016;12:1–23.

Kelly, MB. “Adaptation to Climate Change Through Genetic Accommodation and Assimilation of Plastic Phenotypes.” Phil Trans R Soc B 2019; 374, no. 1768.

Klepsatel P, Gáliková M, De Maio N, Ricci S, Schlötterer C, and Flatt T. Reproductive and post-reproductive life history of wild-caught Drosophila melanogaster under laboratory conditions. J Evol Biol. Switzerland; 2013;26:1508–20.

Kofler R, Orozco-terWengel P, De Maio N, Pandey RV, Nolte V, Futschik A, Kosiol C, and Schlötterer C. PoPoolation: a toolbox for population genetic analysis of next generation sequencing data from pooled individuals. PloS one. Public Library of Science; 2011;6:e15925.

Lande R. Adaptation to an extraordinary environment by evolution of phenotypic plasticity and genetic assimilation. J Evol Biol. Switzerland; 2009;22:1435–46.

Levine MT, Eckert ML, and Begun DJ. Whole-genome expression plasticity across tropical and temperate Drosophila melanogaster populations from Eastern Australia. Mol Biol Evol. United States; 2011;28:249–56.

Levis NA, and Pfennig DW. Evaluating ‘Plasticity-First’ Evolution in Nature: Key Criteria and Empirical Approaches. Trends Ecol Evol. England; 2016;31:563–574.

Levis NA, Isdaner AJ, and Pfennig DW. Morphological novelty emerges from pre-existing phenotypic plasticity. Nat Ecol Evol. England; 2018;2:1289–1297.

Liao Y, Smyth GK, and Shi W. The Subread aligner: fast, accurate and scalable read mapping by seed-and-vote. Nucleic acids research. Oxford Univ Press; 2013;41:e108–e108.

Machado HE, Bergland AO, O’Brien KR, Behrman EL, Schmidt PS, and Petrov DA. Comparative population genomics of latitudinal variation in Drosophila simulans and Drosophila melanogaster. Molecular Ecology. England; 2016;25:723–40.

Mallard F, Nolte V, Tobler R, Kapun M, and Schlötterer C. A simple genetic basis of adaptation to a novel thermal environment results in complex metabolic rewiring in Drosophila. Genome Biol. England; 2018;19:119.

Manenti T, Loeschcke V, Moghadam NN, and Sørensen JG. Phenotypic plasticity is not affected by experimental evolution in constant, predictable or unpredictable fluctuating thermal environments. J Evol Biol. Switzerland; 2015;28:2078–87.

Merilä J, and Hendry AP. Climate change, adaptation, and phenotypic plasticity: the problem and the evidence. Evol Appl. England; 2014;7:1–14.

Mitchell HK, and Lipps LS. Heat shock and phenocopy induction in Drosophila. Cell. United States; 1978;15:907–18.

Montgomery, S. H., & Mank, J. E. (2016). Inferring regulatory change from gene expression: the confounding effects of tissue scaling. Molecular ecology, 25(20), 5114–5128.

Moussian, B., Schwarz, H., Bartoszewski, S., & Nüsslein-Volhard, C. (2005). Involvement of chitin in exoskeleton morphogenesis in *Drosophila melanogaster*. Journal of morphology, 264(1), 117–130.

Nouhaud, P., Tobler, R., Nolte, V., & Schlötterer, C. (2016). Ancestral population reconstitution from isofemale lines as a tool for experimental evolution. Ecology and evolution, 6(20), 7169–7175.

Nunney L. Adapting to a Changing Environment: Modeling the Interaction of Directional Selection and Plasticity. Journal of Heredity. 2016;107:15–24. http://dx.doi.org/10.1093/jhered/esv084

Nussey DH, Postma E, Gienapp P, and Visser ME. Selection on heritable phenotypic plasticity in a wild bird population. Science. 2005;310:304–6.

Palmieri N, Nolte V, Chen J, and Schlötterer C. Genome assembly and annotation of a Drosophila simulans strain from Madagascar. Molecular ecology resources. Wiley Online Library; 2014;.

Pigliucci M. Evolution of phenotypic plasticity: where are we going now? Trends Ecol Evol. England; 2005;20:481–6.

Pigliucci M. Phenotypic plasticity: beyond nature and nurture. Phenotypic plasticity: beyond nature and nurture. Baltimore: Johns Hopkins University Press; 2001.

Porcelli D, Westram AM, Pascual M, Gaston KJ, Butlin RK, and Snook RR. Gene expression clines reveal local adaptation and associated trade-offs at a continental scale. Scientific Reports. England; 2016;6:32975.

Price TD, Qvarnström A, and Irwin DE. The role of phenotypic plasticity in driving genetic evolution. Proceedings of the Royal Society of London Series B: Biological Sciences. 2003;270:1433–1440.

Robinson MD, McCarthy DJ, and Smyth GK. edgeR: a Bioconductor package for differential expression analysis of digital gene expression data. Bioinformatics. Oxford Univ Press; 2010;26:139–140.

Sgrò CM, Terblanche JS, and Hoffmann AA. What Can Plasticity Contribute to Insect Responses to Climate Change? Annu Rev Entomol. United States; 2016;61:433–51.

Sørensen JG, Kristensen TN, and Overgaard J. Evolutionary and ecological patterns of thermal acclimation capacity in Drosophila: is it important for keeping up with climate change? Curr Opin Insect Sci. Netherlands; 2016;17:98–104.

Suzuki Y, and Nijhout HF. Evolution of a Polyphenism by Genetic Accommodation. Science. 2006;311:650–652.

Team RC. R: A language and environment for statistical computing. Vienna: R Foundation for Statistical Computing; 2016.

Vázquez DP, Gianoli E, Morris WF, and Bozinovic F. Ecological and evolutionary impacts of changing climatic variability. Biol Rev Camb Philos Soc. England; 2017;92:22–42.

Via S, Gomulkiewicz R, De Jong G, Scheiner SM, Schlichting CD, and Van Tienderen PH. Adaptive phenotypic plasticity: consensus and controversy. Trends Ecol Evol. England; 1995;10:212–7.

Via, S. (1993). Adaptive phenotypic plasticity: target or by-product of selection in a variable environment?. The American Naturalist, 142(2), 352–365.

Wang L, Wang S, and Li W. RSeQC: quality control of RNA-seq experiments. Bioinformatics. Oxford Univ Press; 2012;28:2184–2185

West-Eberhard MJ. Developmental plasticity and evolution. Developmental plasticity and evolution. Oxford; New York: Oxford University Press; 2003.

Whitman DW, and Ananthakrishna TN. Phenotypic plasticity of insects: mechanisms and consequences. Phenotypic plasticity of insects: mechanisms and consequences. USA: Science Publishers; 2009.

Wu TD, and Nacu S. Fast and SNP-tolerant detection of complex variants and splicing in short reads. Bioinformatics. Oxford Univ Press; 2010;26:873–881.

Yampolsky LY, Glazko GV, and Fry JD. Evolution of gene expression and expression plasticity in long-term experimental populations of Drosophila melanogaster maintained under constant and variable ethanol stress. Mol Ecol. England; 2012;21:4287–99.

Zhao L, Wit J, Svetec N, and Begun DJ. Parallel Gene Expression Differences between Low and High Latitude Populations of Drosophila melanogaster and *D. simulans*. PLoS Genetics. United States; 2015;11:e1005184.

Zhou S, Campbell TG, Stone EA, Mackay TF, and Anholt RR. Phenotypic plasticity of the Drosophila transcriptome. PLoS Genet United States; 2012;8:e1002593.

