## Supplementary figures and images for "The evolution of phenotypic plasticity in response to temperature stress"

### Suplemental Figure

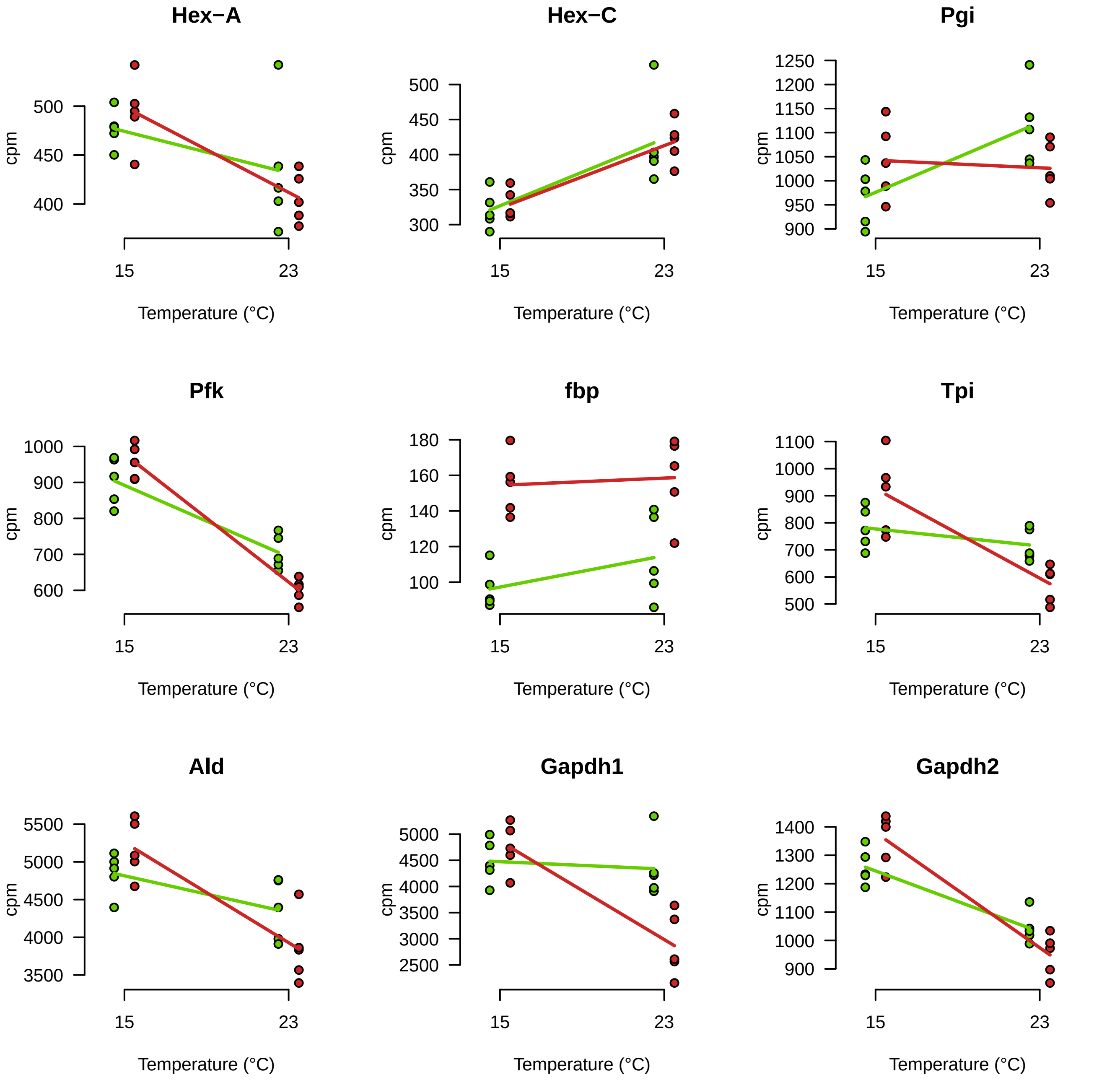

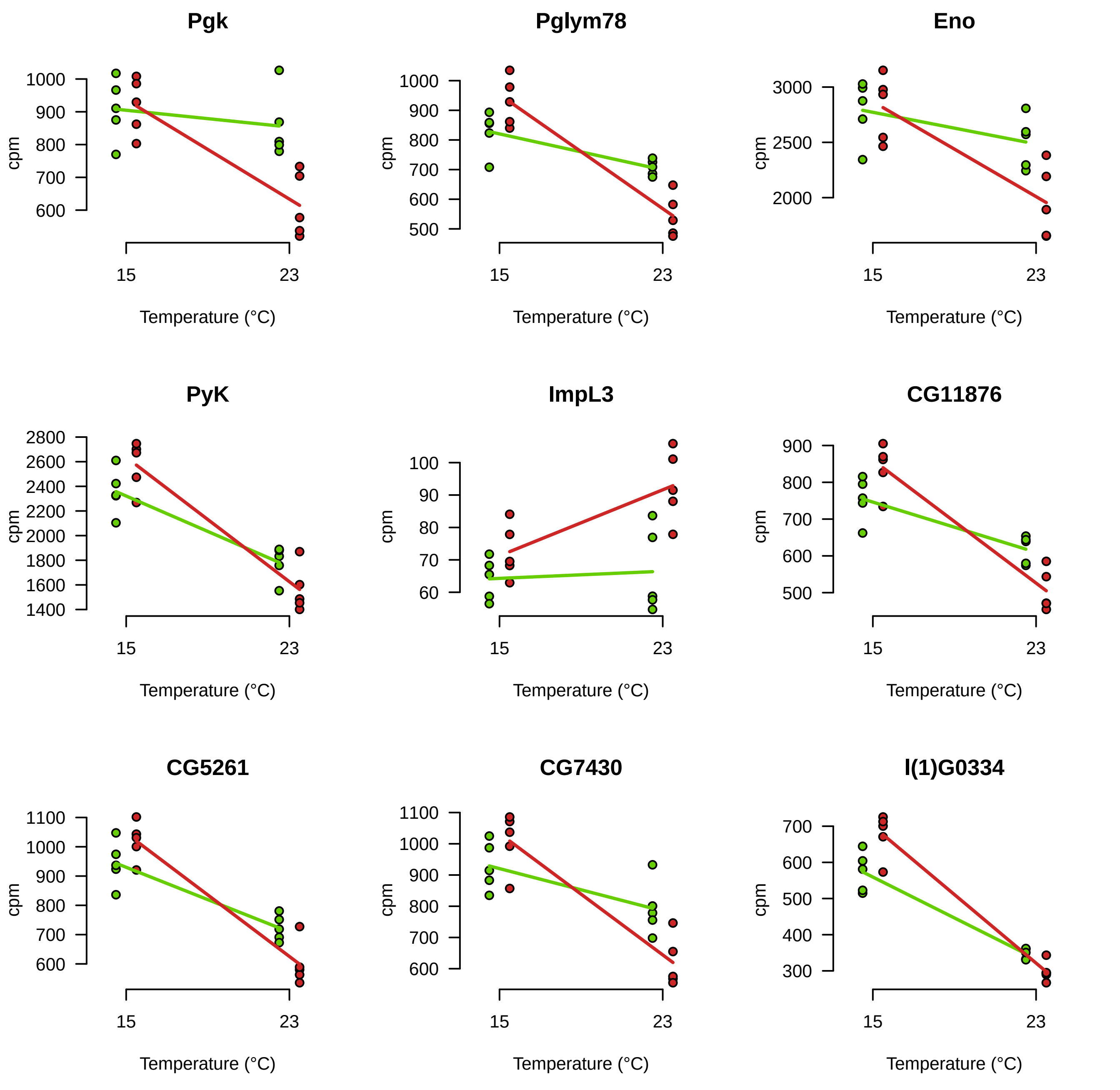
