## Supplementary material for "The evolution of phenotypic plasticity in response to temperature stress": Suplemental Figure

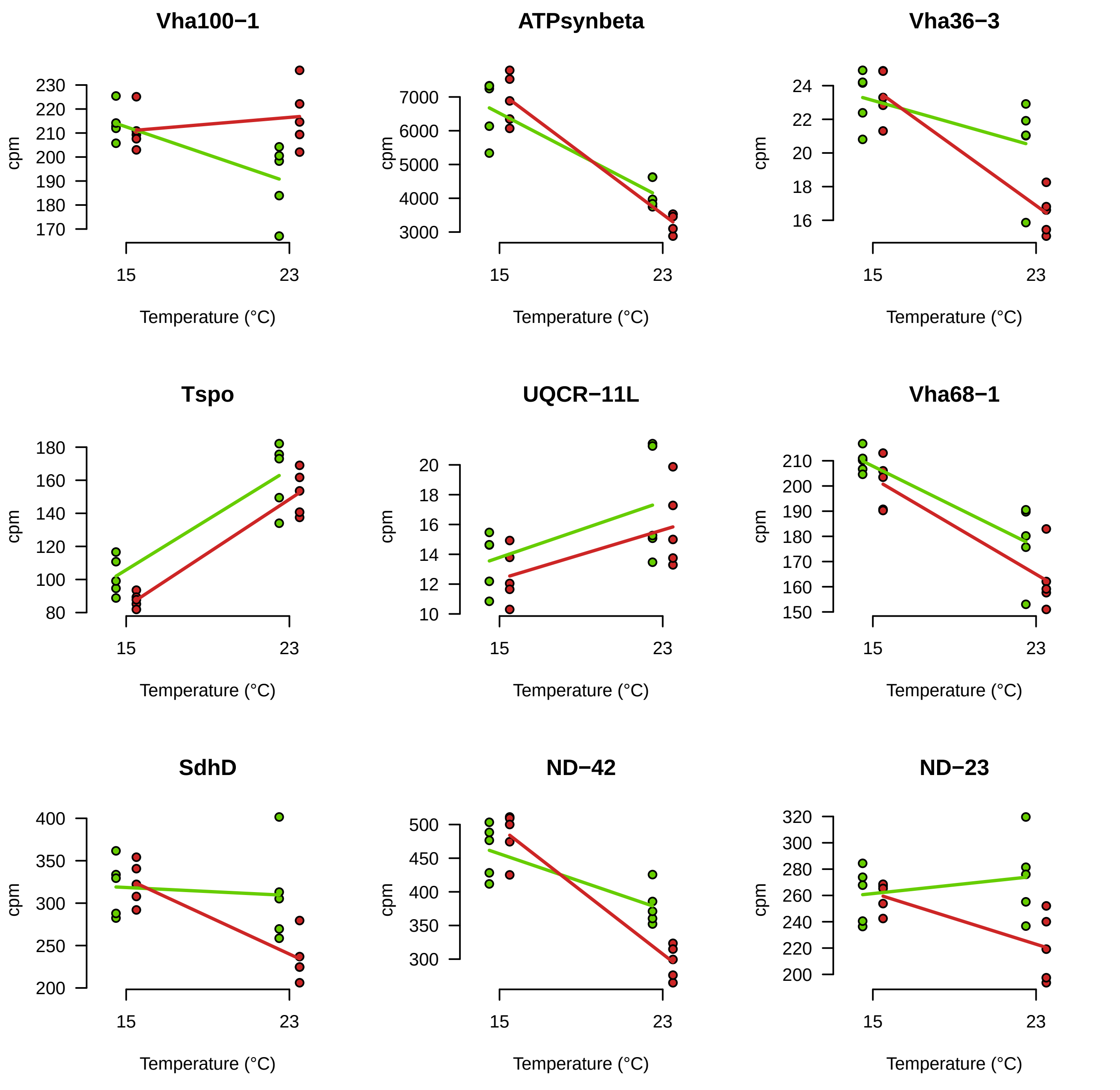

Vha100−1
ATPsynbeta
Vha36−3
Tspo
UQCR−11L
Vha68−1
SdhD
ND−42
ND−23
cpm
Temperature (°C)
15
23

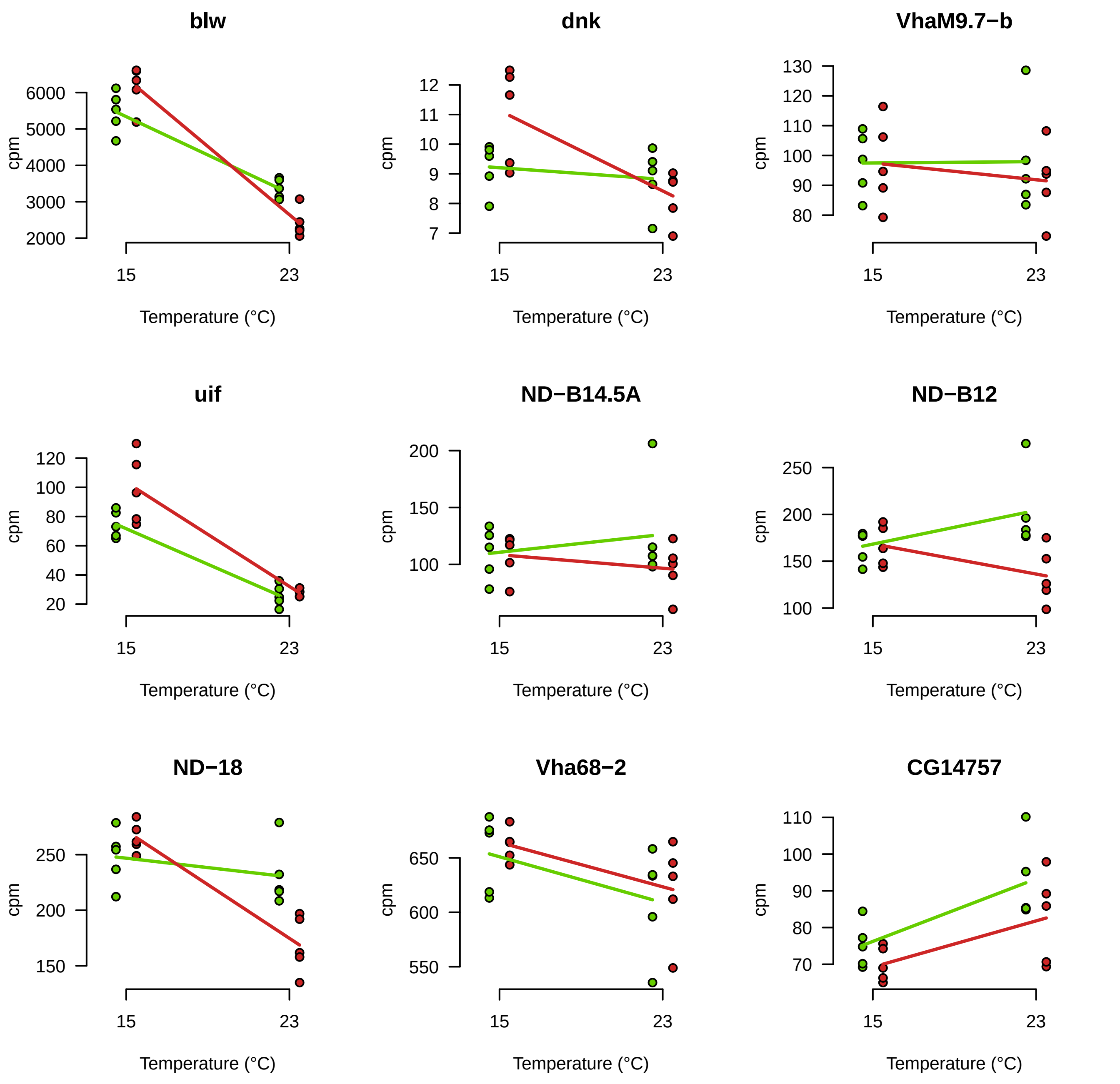
blw
dnk
VhaM9.7−b
uif
ND−B14.5A
ND−B12
ND−18
Vha68−2
CG14757
cpm
Temperature (°C)
15
23

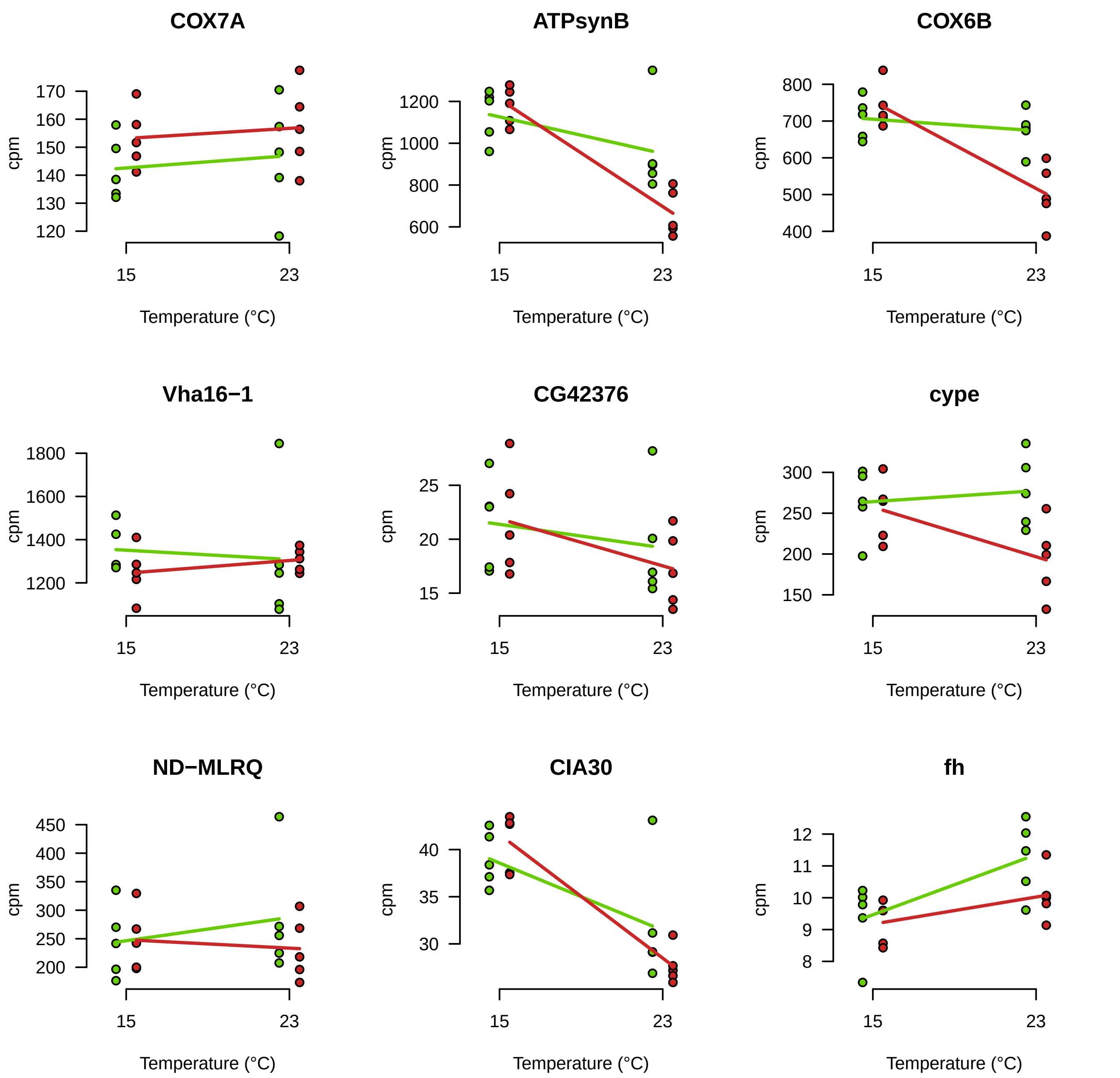

COX7A
ATPsynB
COX6B
Vha16−1
CG42376
cype
ND−MLRQ
CIA30
fh
cpm
Temperature (°C)
15
23

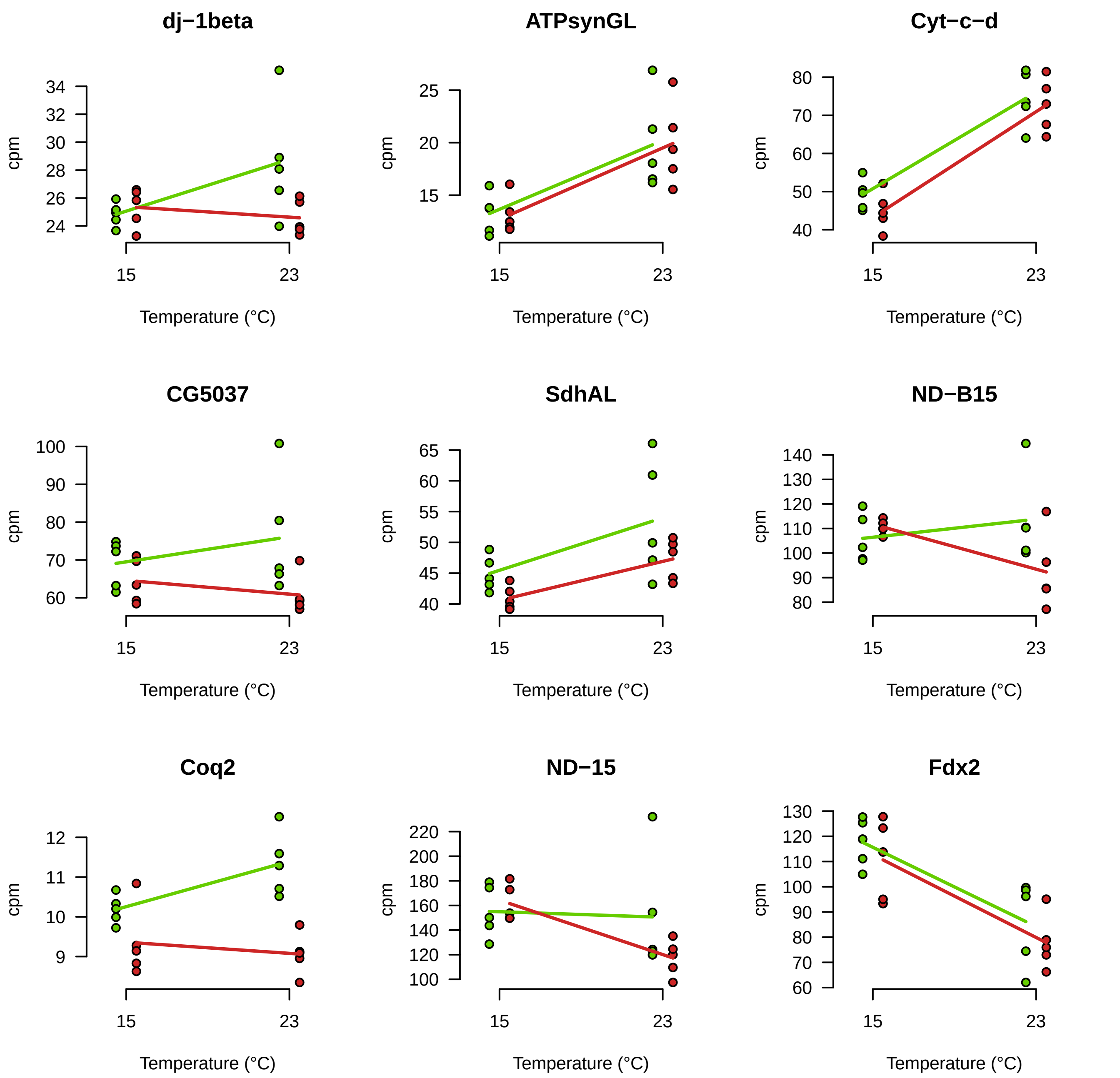

dj−1beta
ATPsynGL
Cyt−c−d
CG5037
SdhAL
ND−B15
Coq2
ND−15
Fdx2
cpm
Temperature (°C)
15
23

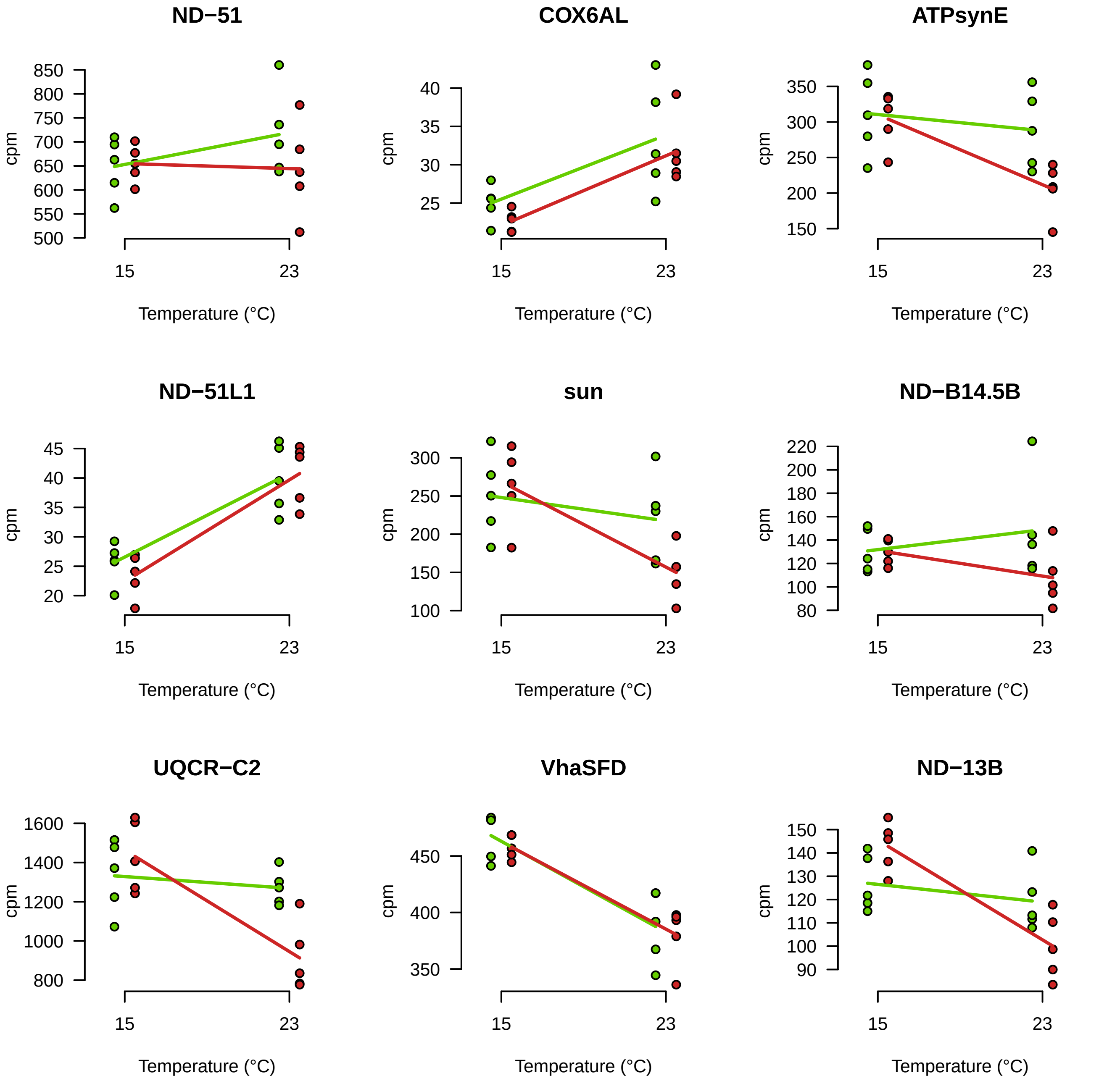

ND−51
COX6AL
ATPsynE
ND−51L1
sun
ND−B14.5B
UQCR−C2
VhaSFD
ND−13B
cpm
Temperature (°C)
15
23

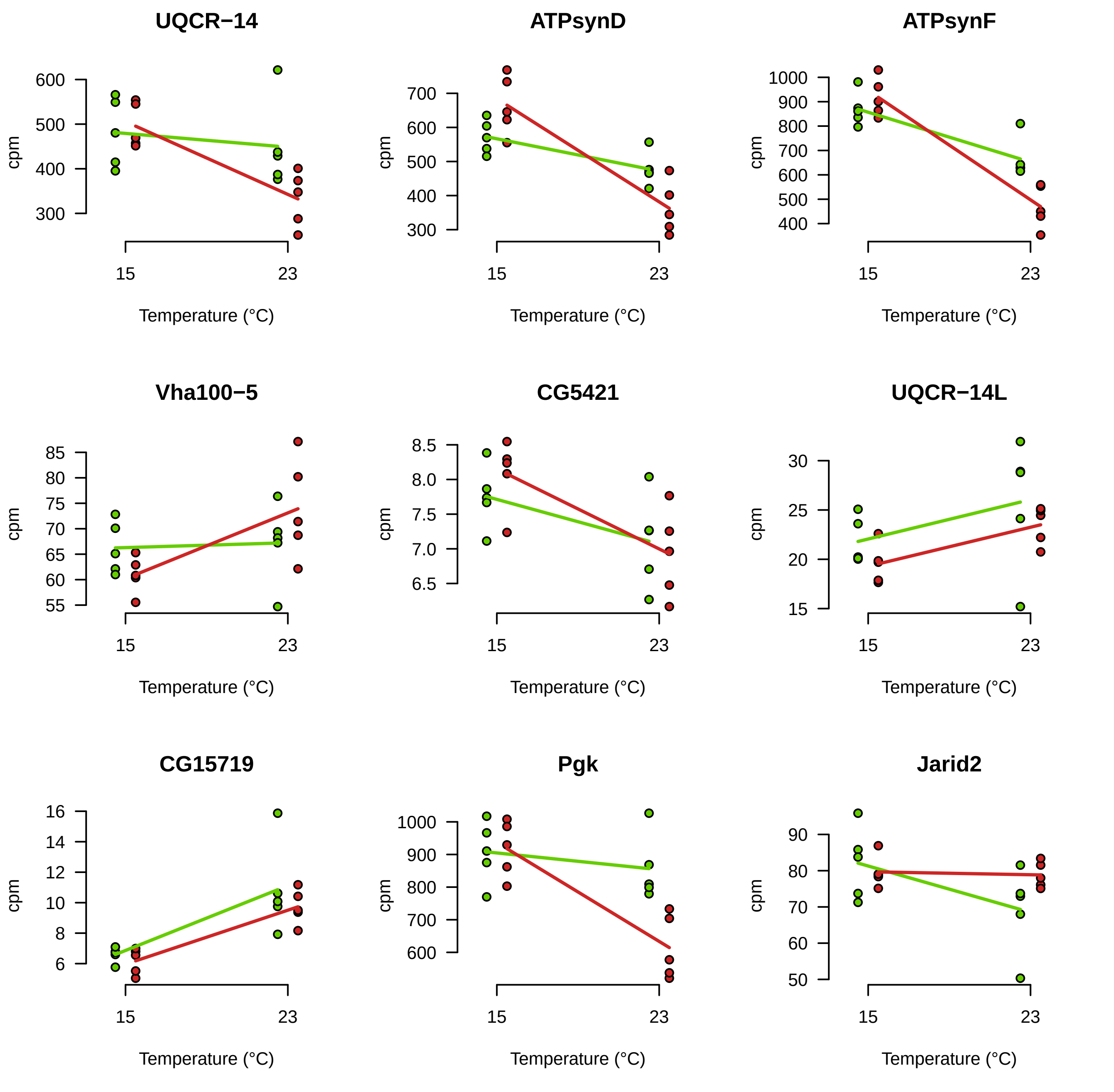
UQCR−14
ATPsynD
ATPsynF
Vha100−5
CG5421
UQCR−14L
CG15719
Pgk
Jarid2
cpm
Temperature (°C)
15
23

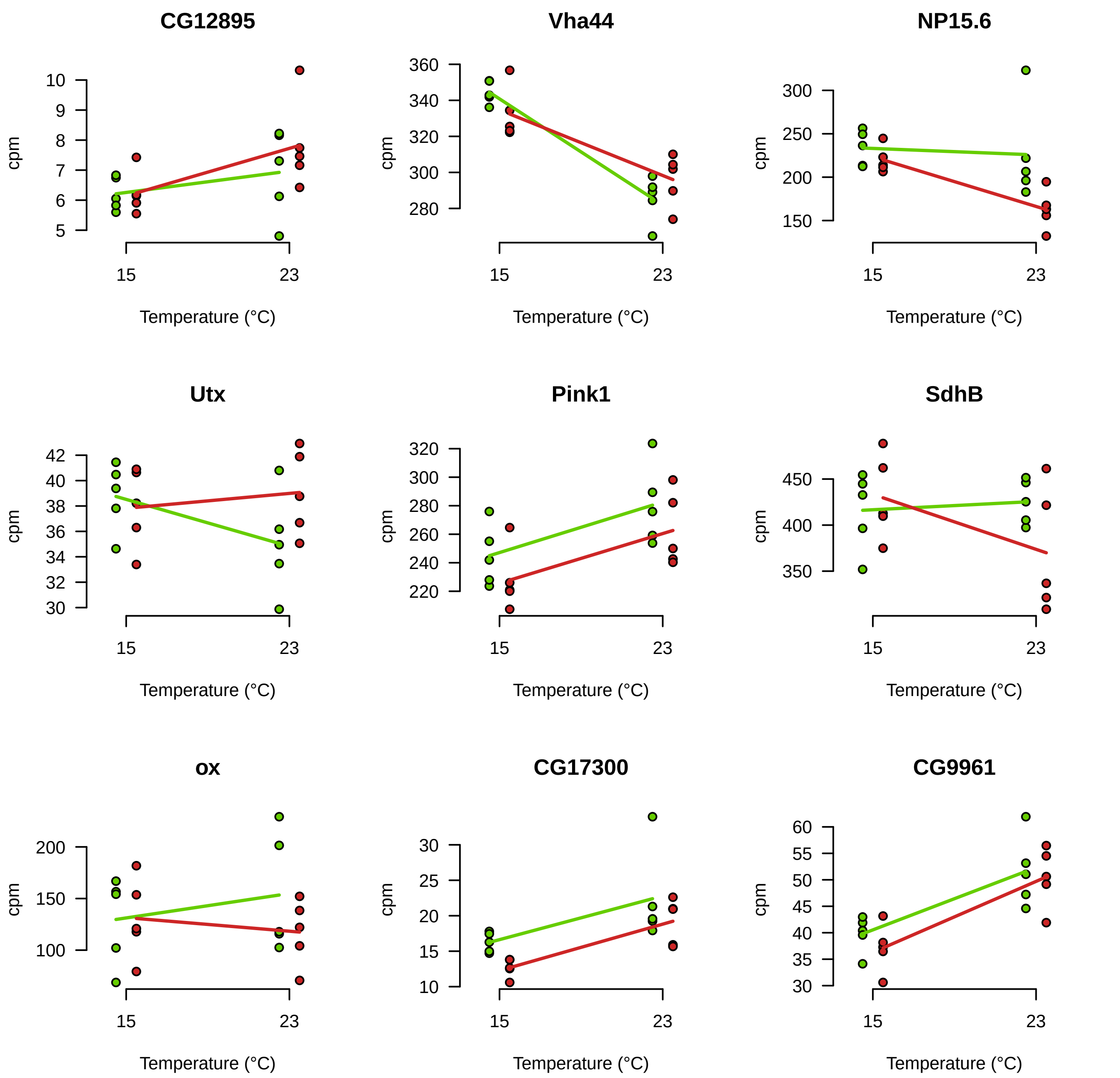

CG12895
Vha44
NP15.6
Utx
Pink1
SdhB
ox
CG17300
CG9961
cpm
Temperature (°C)
15
23

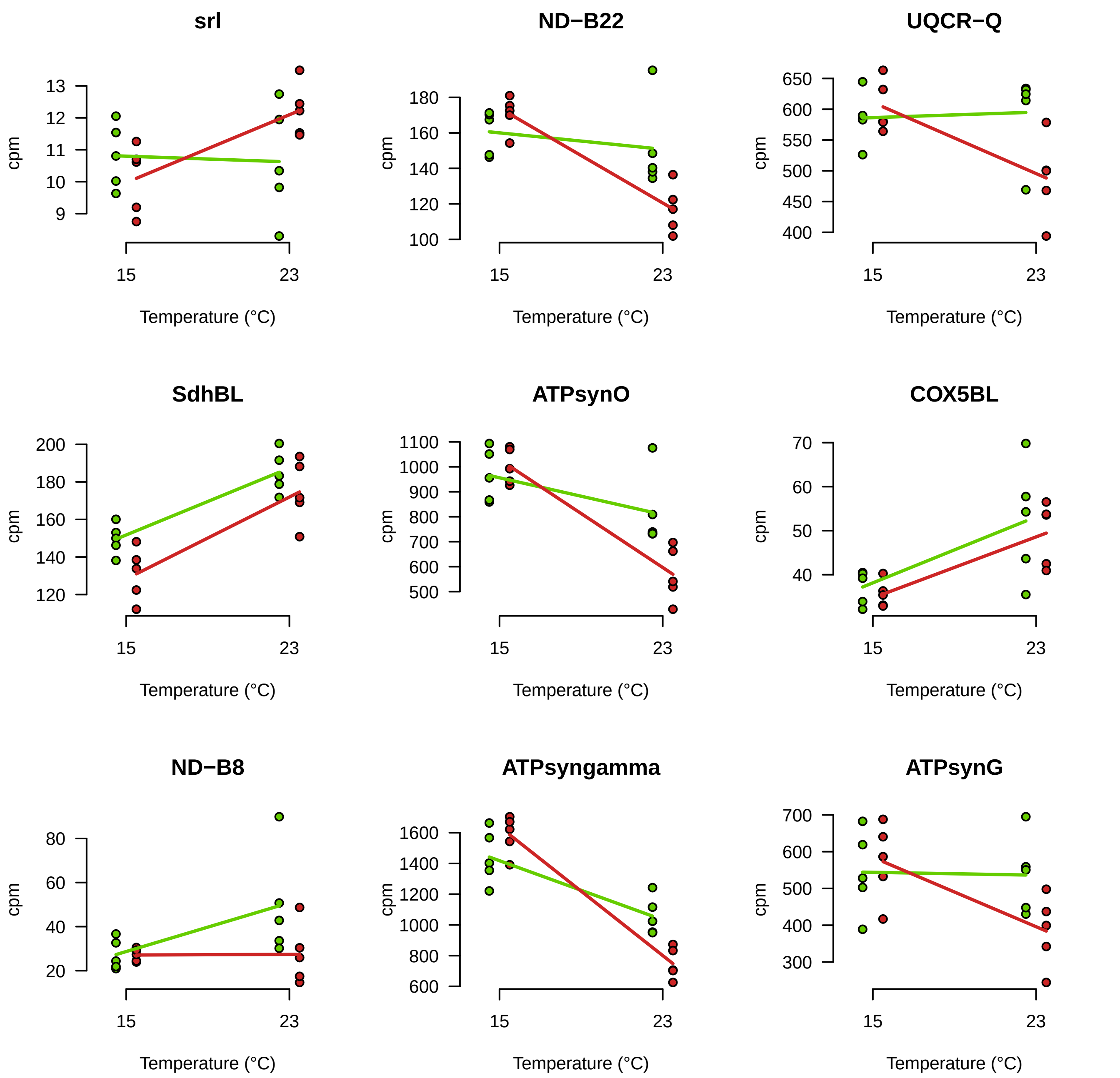
srl
ND−B22
UQCR−Q
SdhBL
ATPsynO
COX5BL
ND−B8
ATPsyngamma
ATPsynG
cpm
Temperature (°C)
15
23

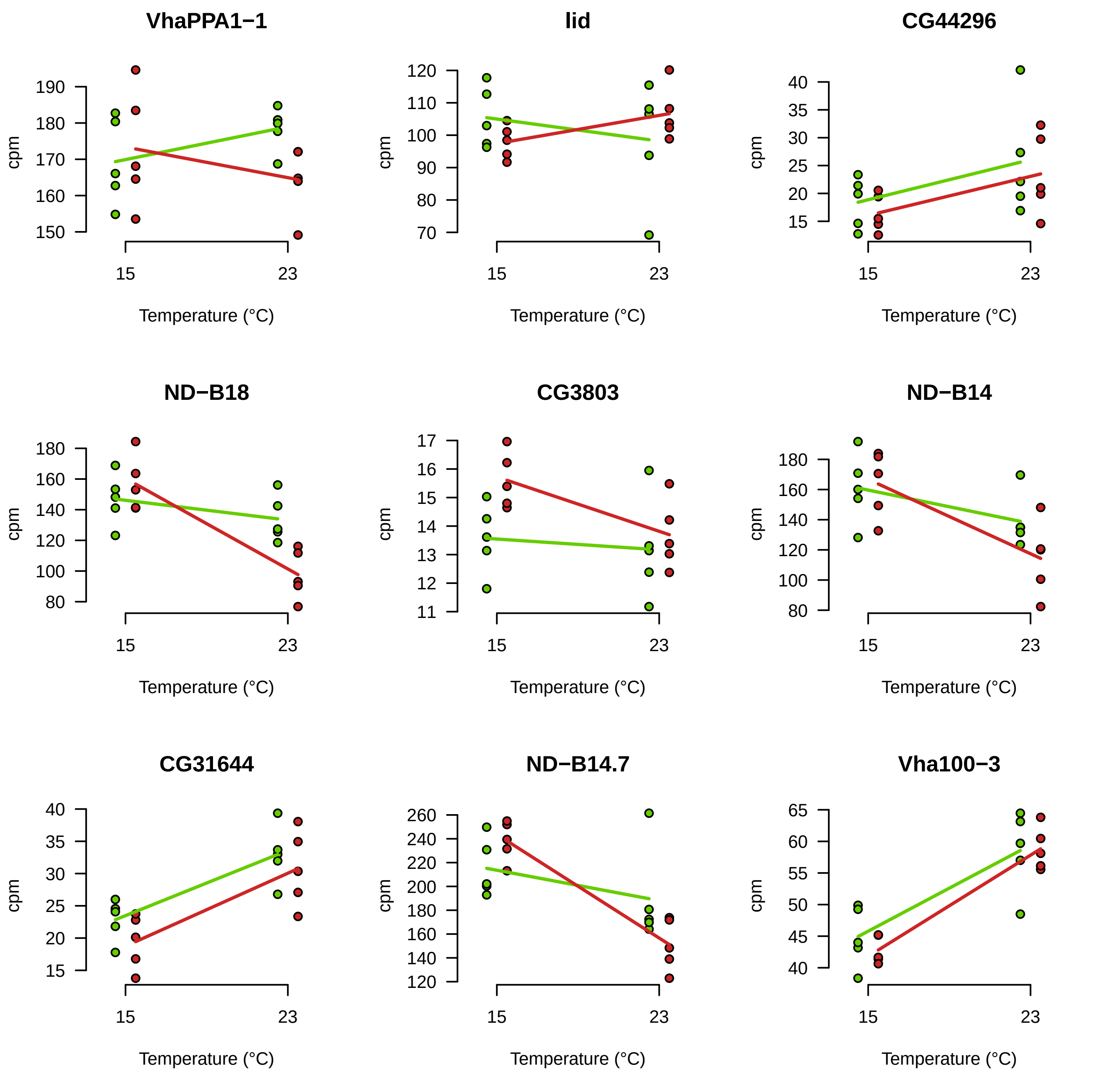

VhaPPA1−1
lid
CG44296
ND−B18
CG3803
ND−B14
CG31644
ND−B14.7
Vha100−3
cpm
Temperature (°C)
15
23

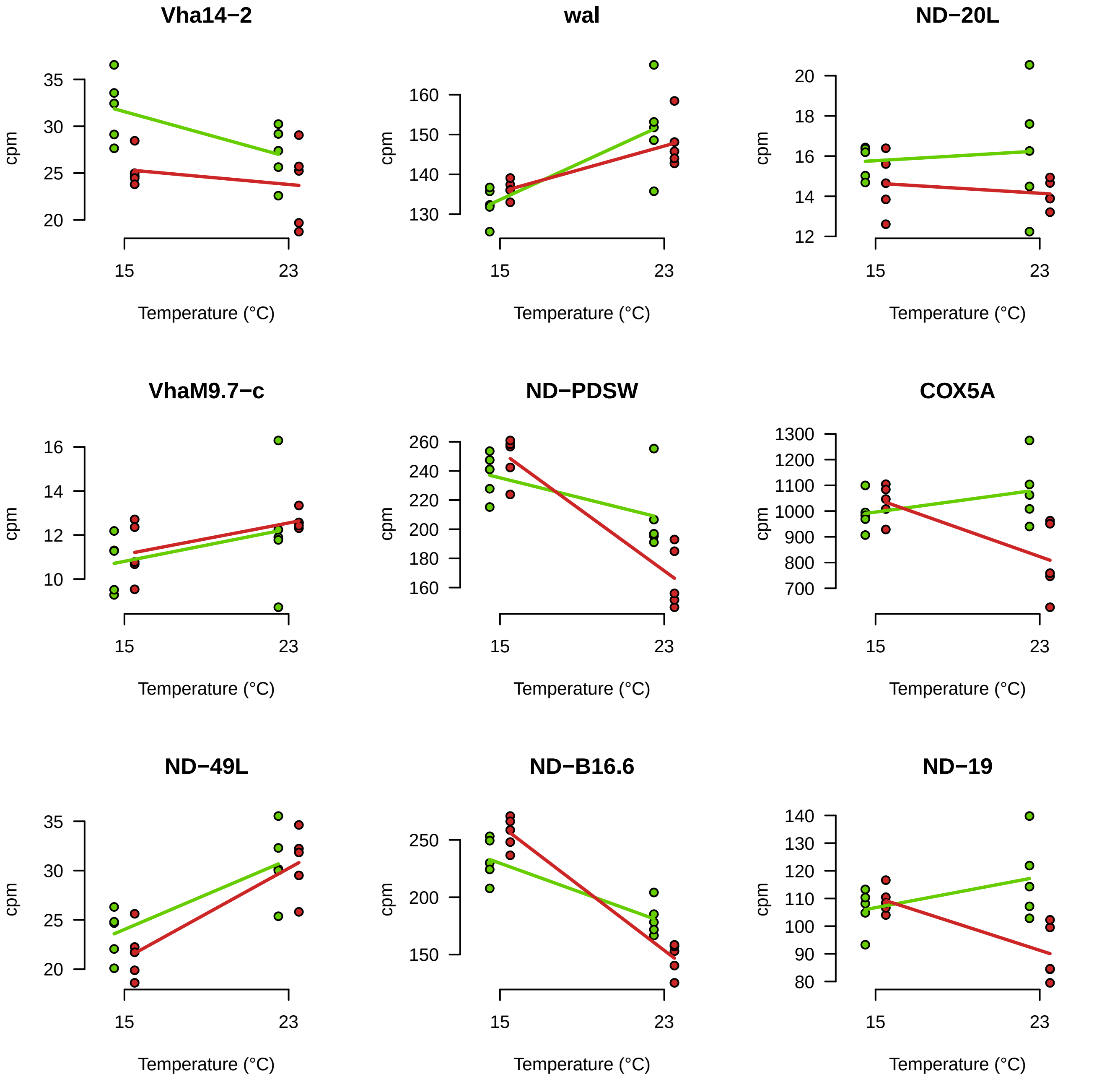
Vha14−2
wal
ND−20L
VhaM9.7−c
ND−PDSW
COX5A
ND−49L
ND−B16.6
ND−19
cpm
Temperature (°C)
15
23
35
30
25
20
160
150
140
130
20
18
16
14
12
10
260
240
220
200
180
1300
1200
1100
1000
900
800
700
250
140
120
110
100
90
80

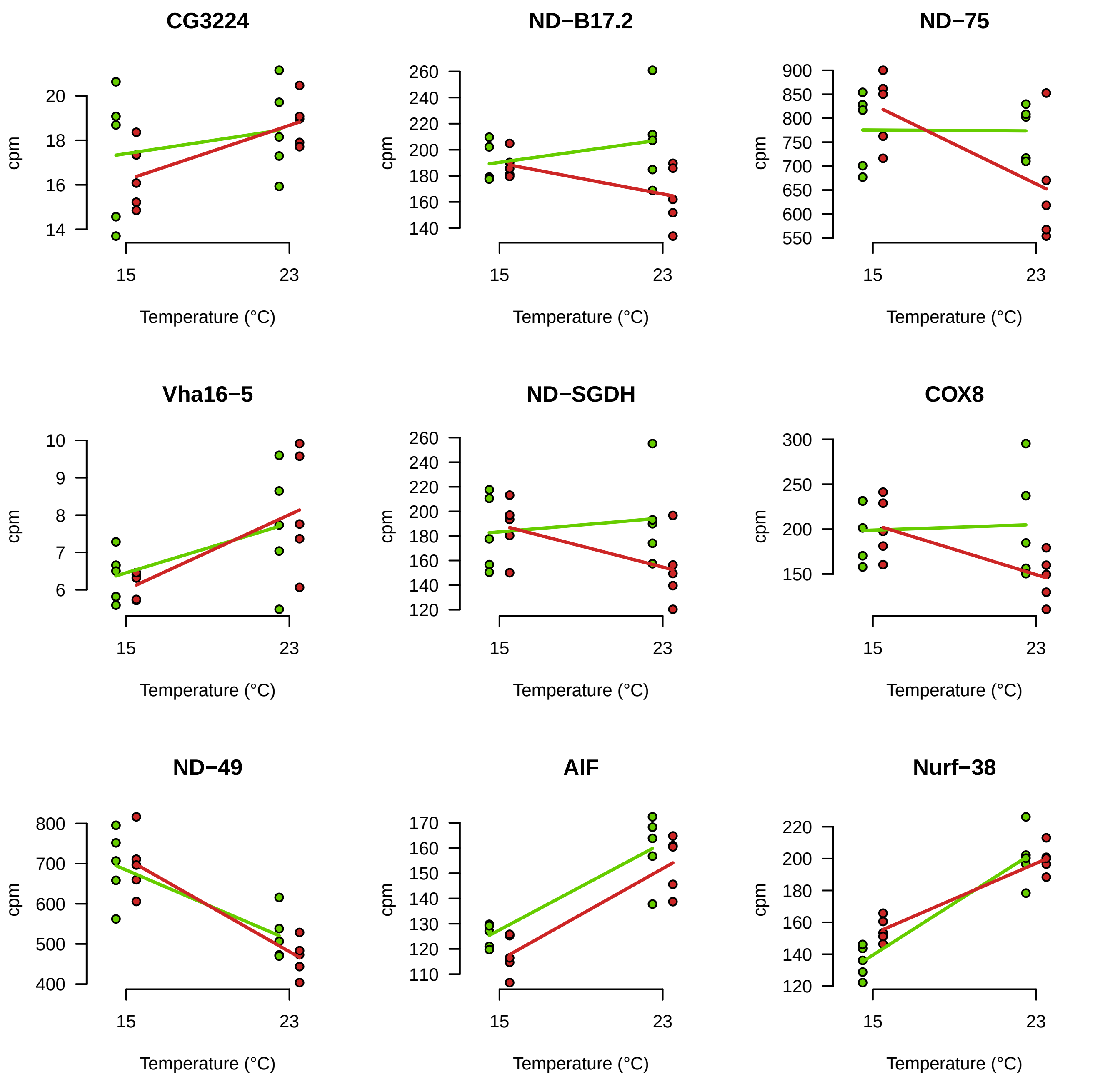
CG3224
ND−B17.2
ND−75
Vha16−5
ND−SGDH
COX8
ND−49
AIF
Nurf−38
cpm
Temperature (°C)
15
23

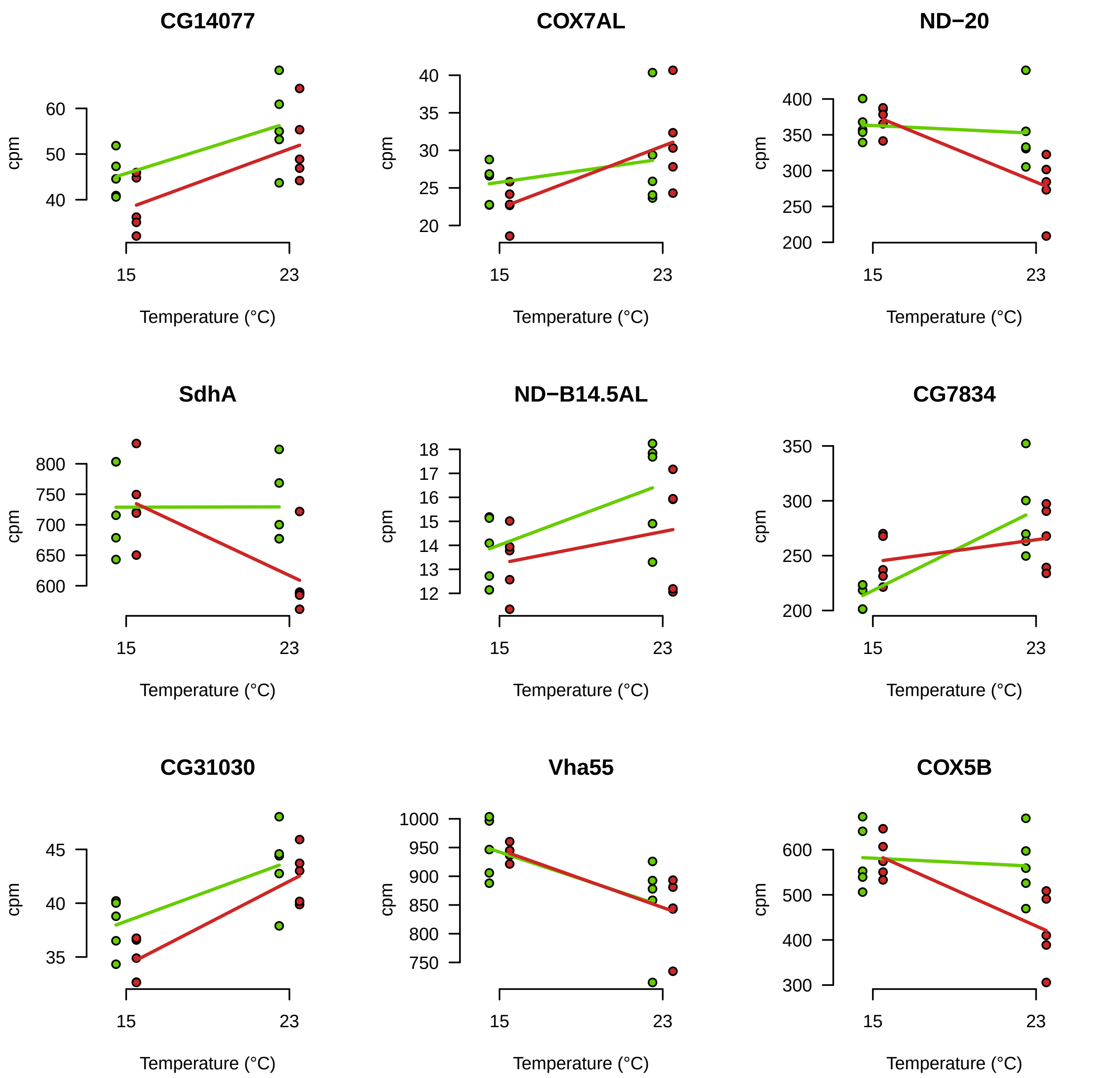

CG14077
COX7AL
ND−20
SdhA
ND−B14.5AL
CG7834
CG31030
Vha55
COX5B
cpm
Temperature (°C)
15
23

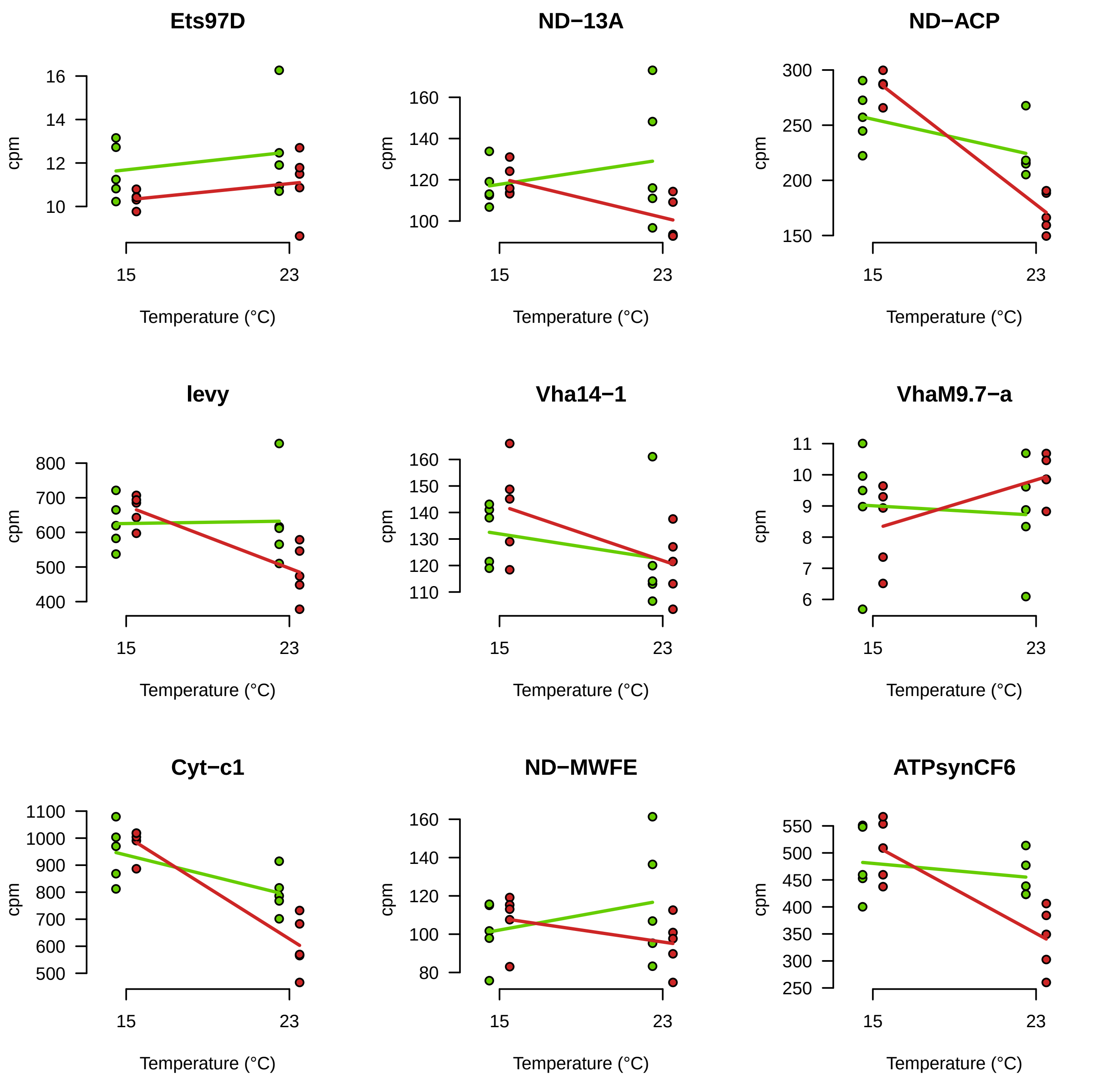
Ets97D
ND−13A
ND−ACP
levy
Vha14−1
VhaM9.7−a
Cyt−c1
ND−MWFE
ATPsynCF6
cpm
Temperature (°C)
15
23

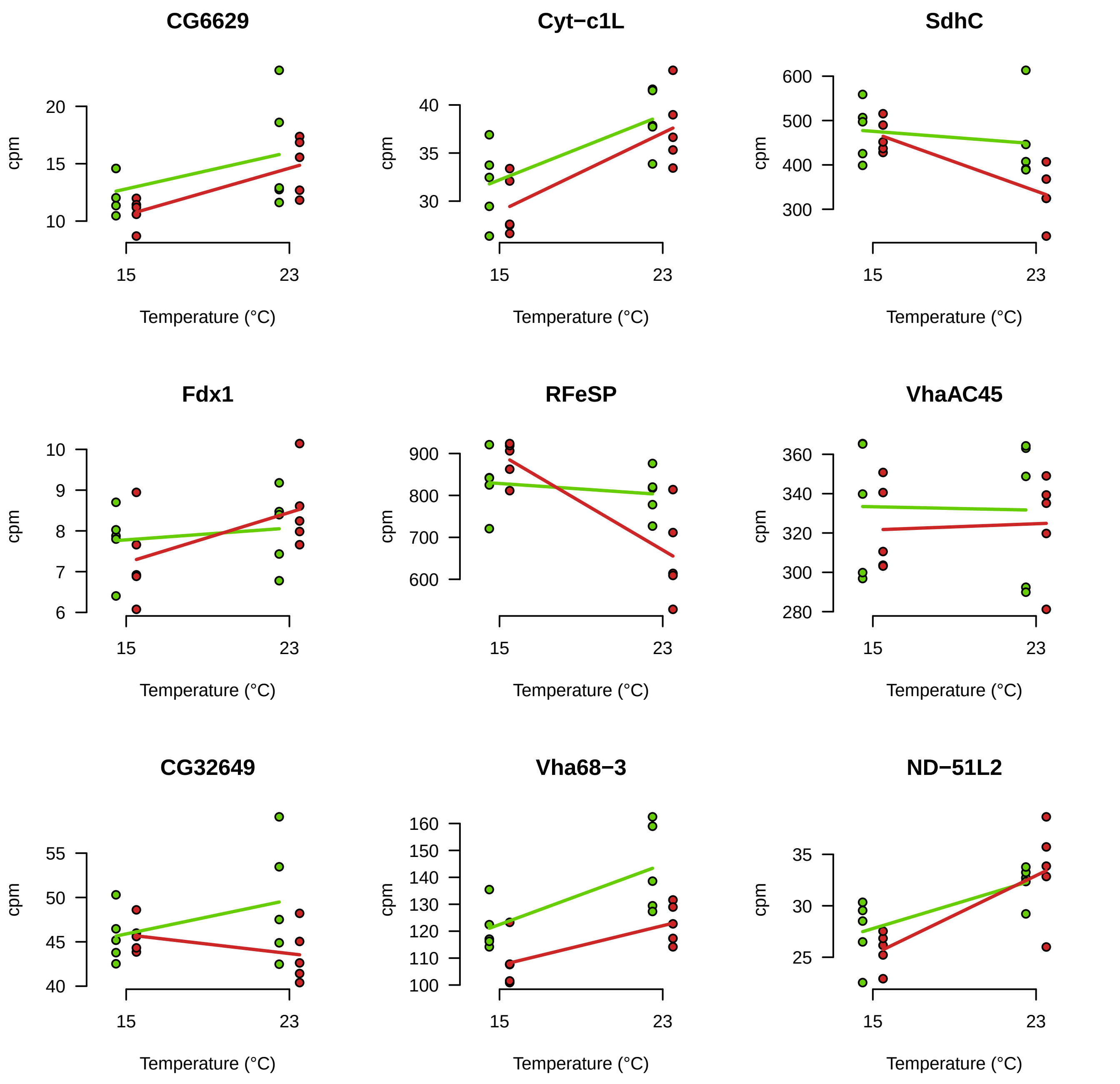

CG6629
Cyt−c1L
SdhC
Fdx1
RFeSP
VhaAC45
CG32649
Vha68−3
ND−51L2
cpm
Temperature (°C)
15
23
10
15
20
30
35
40
300
400
500
600
6
7
8
9
10
600
700
800
900
280
300
320
340
360
40
45
50
55
100
110
120
130
140
150
160
25
30
35

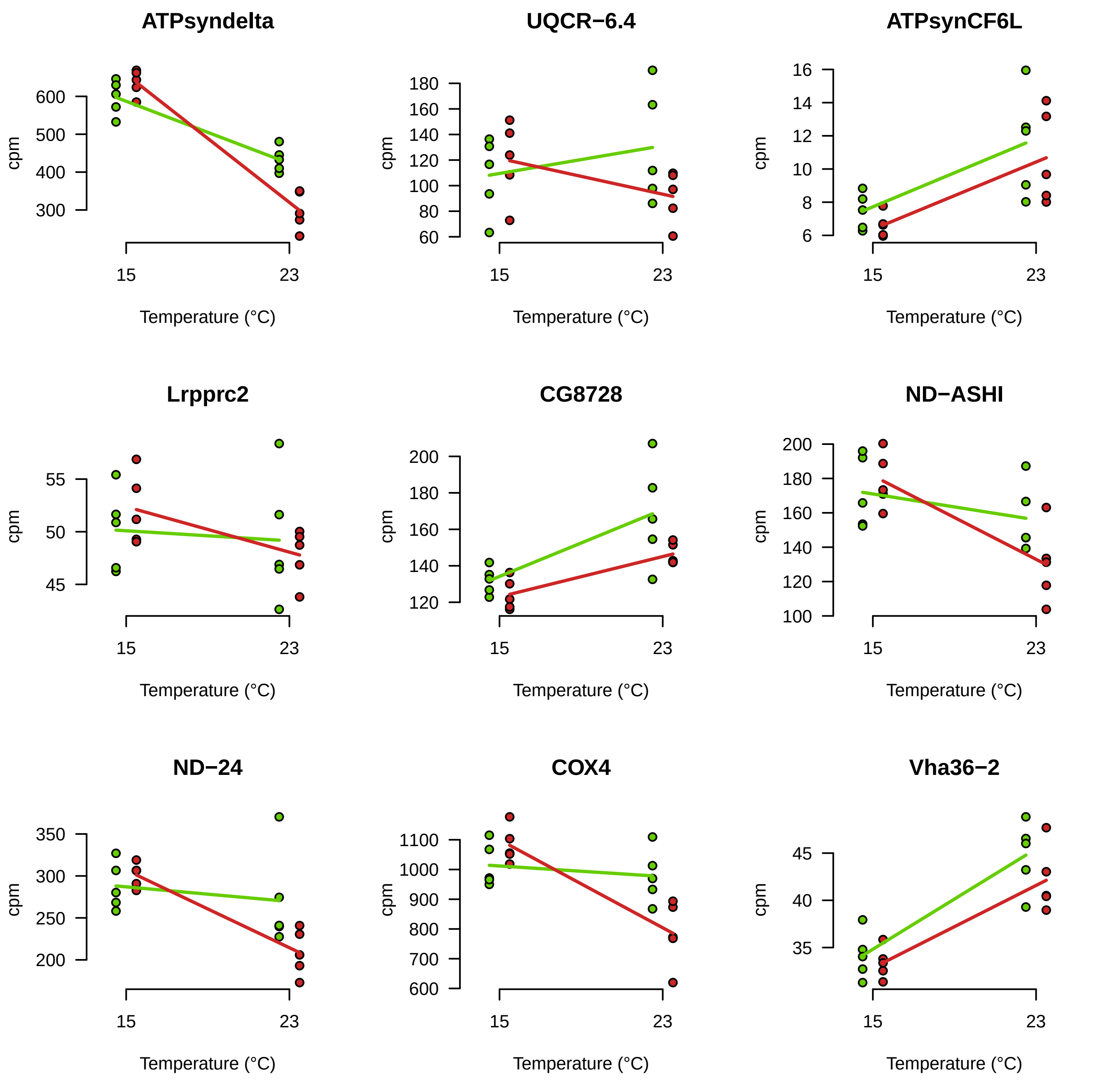
ATPsyndelta
UQCR−6.4
ATPsynCF6L
Lrpprc2
CG8728
ND−ASHI
ND−24
COX4
Vha36−2
cpm
Temperature (°C)
15
23

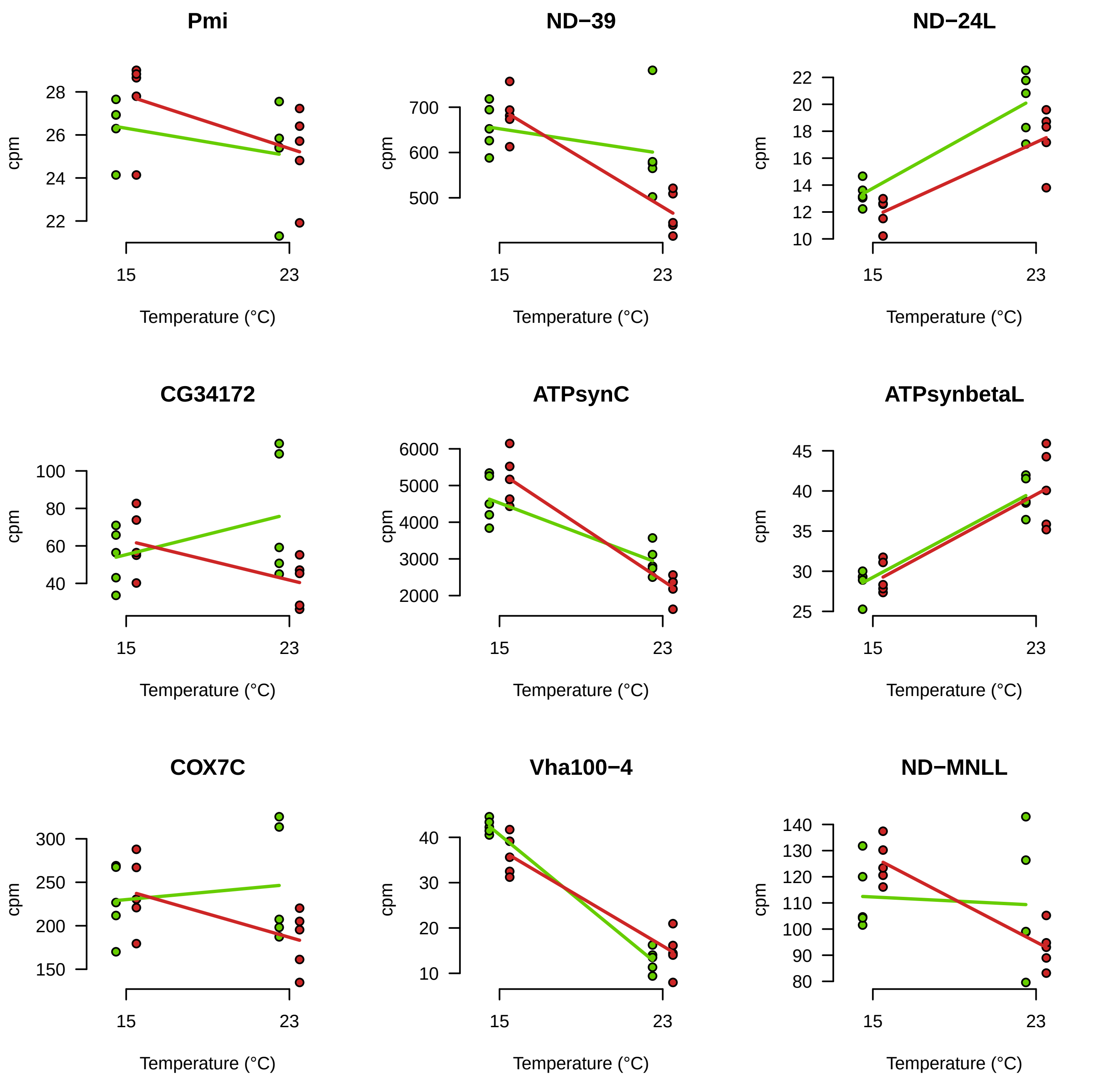

Pmi
ND−39
ND−24L
CG34172
ATPsynC
ATPsynbetaL
COX7C
Vha100−4
ND−MNLL
cpm
Temperature (°C)
15
23

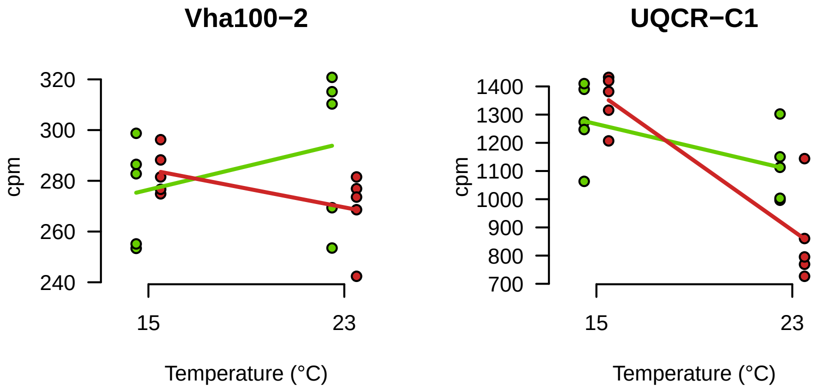
Vha100−2
320
300
280
260
240
cpm
15
23
Temperature (°C)
UQCR−C1
1400
1300
1200
1100
1000
900
800
700
cpm
15
23
Temperature (°C)
